# Dopamine depletion leads to pathological synchronization of distinct basal ganglia loops in the beta band

**DOI:** 10.1101/2022.10.10.511532

**Authors:** Andrea Ortone, Alberto Arturo Vergani, Riccardo Mannella, Alberto Mazzoni

## Abstract

Motor symptoms of Parkinson’s Disease (PD) are associated with dopamine deficits and pathological oscillation of basal ganglia (BG) neurons in the *β* range ([12-30] Hz). However, how the dopamine depletion affects the oscillation dynamics of BG nuclei is still unclear. With a spiking neurons model, we here captured the features of BG nuclei interactions leading to oscillations in dopamine-depleted condition. We found that both the loop between subthalamic nucleus and Globus Pallidus pars externa (GPe) and the loop between striatal fast spiking and medium spiny neurons and GPe displayed resonances in the *β* range, and synchronized to a common *β* frequency through interaction. Crucially, the synchronization depends on dopamine depletion: the two loops were largely independent for high levels of dopamine, but progressively synchronized as dopamine was depleted due to the increased strength of the striatal loop. Our results highlight the role of the interplay between the GPe-STN and the GPe-striatum loop in generating sustained *β* oscillations in PD subjects, and explain how this interplay depends on the level of dopamine. This paves the way to the design of therapies specifically addressing the onset of pathological *β* oscillations.

**Author summary:** Parkinson’s Disease is associated to the death of neurons generating a particular neurotransmitter: the dopamine. Motor symptoms of PD, on the other hand, are known to be due to dysfunctions in a particular subcortical area of the brain, the BG network. In particular, the BG network develops pathological oscillations in a specific frequency range (*β*: [12-30] Hz). What is unclear is how dopamine depletion leads to these oscillations. In this work we developed a BG network model and we found the actual reason for these abnormal oscillations is the synchronization of two loops within the network that are individually oscillating in the *β* range. For healthy level of dopamine the two loops are decoupled and the oscillation power is low. When dopamine is depleted (as in PD) the two loops synchronize and originate the pathological oscillations associated with motor symptoms.

## Introduction

The basic architecture of the basal ganglia (BG) network (see Fig 1) consists of the striatum (STR), the globus pallidus, divided into pars interna (GPi) and pars esterna (GPe), the substantia nigra (reticulata SNr and compacta SNc) and the subthalamic nucleus (STN) [1, 2]. In the striatum there are three interacting populations of neurons: D1 neurons excited by dopamine, D2 neurons inhibited by dopamine and Fast Spiking Neurons (FSN).

**Fig 1.**
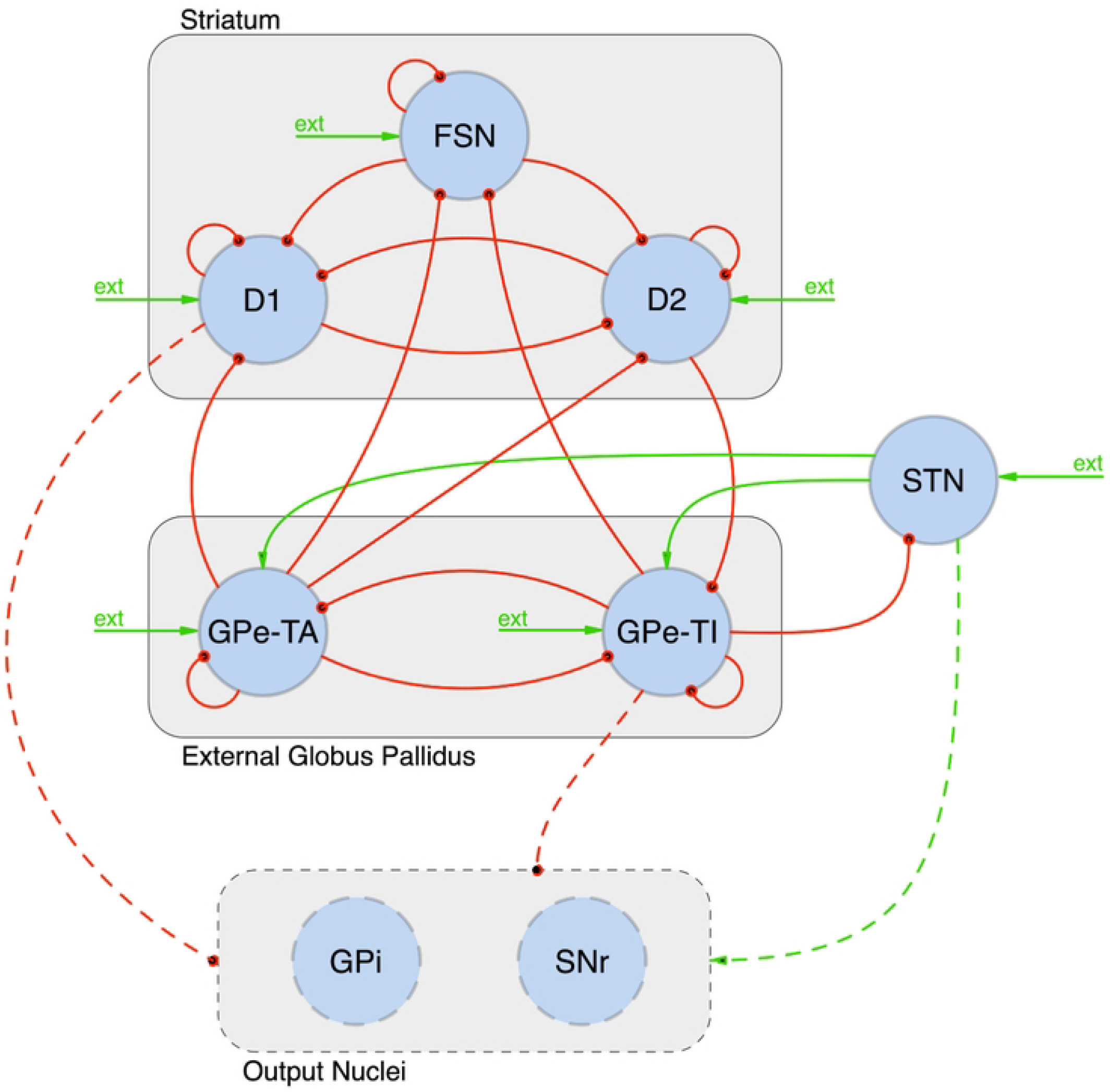
Architecture of the adopted Complete Model of the basal ganglia. FSN, D1 and D2: striatal Fast Spike Neurons, and medium spiny neurons with D1 and D2 dopamine receptors; GPe-TA and GPe-TI: globus pallidus externa type A and type I; STN: subthalamic nucleus; GPi: globus pallidus interna; SNr: substantia nigra pars reticulata; ext: external poissonian input. By convention red/green arrows are inhibitory/excitatory projections. Dashed elements have not been included in our model.

Parkinson’s Disease (PD) is a common neurodegenerative disease affecting about 0.3% of the world’s population [3]. Etiologically, PD follows the progressive death of dopaminergic neurons in the substantia nigra. The consequent condition of dopamine depletion (*D*_*d*_) leads to an alteration of the balance between D1 excitation and D2 inhibition [4] and this reverberates over the whole network. The main motor symptoms of PD (akinesia [5], bradykinesia [6], tremor [7], freezing of gait [8, 9]) correlate with this dopamine deficiency.

Following the condition of dopamine depletion and the consequent alteration of the striatal activity (higher spiking rate in D2 and lower in D1) [10], pathological *β* oscillations [12 − 30] Hz emerge in the striatum [11–14], in GPi and GPe [15, 16], and in the STN [17, 18]. Experimental recordings ([19, 20]) highlighted that such pathological activity is not characterized by constant intensity, but consists of phasic bursts. Despite the wealth of studies in the subject, the origin of these *β* activities is still debated.

Some hypotheses argue that the oscillations depend on the interaction between GPe and STN [21]. In support of this hypothesis, it is known that the architecture of the STN-GPe circuit is prone to generate oscillatory behavior [22]: STN is a glutamatergic nucleus projecting substantially to GPe and GPe is a GABAergic nucleus projecting feedback to STN [23]. Moreover, in PD, these nuclei present prominent and coherent oscillations in the firing rates [22, 24] and are effective targets of Deep Brain Stimulation (DBS) therapy [25–27]. However, whether *β* band activity can be produced in the STN–GPe circuit is still debated [28]. Several computational models investigated STN-GPe *β* band loop: Kumar et al [29] showed that *β* oscillations could emerge from the STN-GPe network as a consequence of increased inhibitory input from the D2 population in the striatum. Similar results were obtained by Gillies at al [30] and further explored in [31–34].

An alternative explanation for the experimental observations is that the cortex might be a source of *β* oscillations [35, 36]. In accordance with this hypothesis, Van Albada et al. [37] suggested a cortico-thalamic loop source of *β* oscillations, which spread into the basal ganglia.

The third hypothesis is the one of a prominent role of the interaction between GPe and STR: experimental recordings in rats [38–42] and mice [43–46] suggest that the GPe nucleus and the pallido-striatal pathway could play a major role. In vivo observations of pauses in fast spiking neurons FSN support this hypothesis as transient drop in the activity might arise from interactions with GPe [47] (see also [48, 49] for an extended review). From a computational point of view, a model of the GPe-striatal circuit has been developed by Corbit et al [50], who investigated the onset of pathological *β* oscillations in a subnetwork consisting by the inhibitory synapses FSN → D2-MSN → GPe.

In this work we propose a comprehensive model including all the loops with the aim of addressing the aforementioned hypothesis on the mechanisms underlying abnormal *β* oscillations in PD. In line with this purpose, the starting point of our work is the network developed by Lindahl and Kotalewski [51] which includes all major nuclei and connections of the BG. In this context, we identify the major sources of *β* activity, investigate the role of their interaction and highlight the effects of dopamine depletion.

## Materials and methods

### 1.1 Basal ganglia network model

Our model of the basal ganglia (Fig 1) includes:

- three striatal populations: D1-type dopamine receptor Medium Spiny Neurons (D1); D2-type dopamine receptor Medium Spiny Neurons (D2) and a population of Fast Spiking (inter-)Neurons (FSN);
- the external Globus Pallidus divided into two populations that will be labelled as GPe-TA (characterized by a lower discharge rate and by a negligible input from striatal populations) and GPe-TI (with a higher activity and receiving input from D2);
- the SubThalamic Nucleus (STN).

Each population presents specific size (see Table 2) and neuron model (see section 1.2). Single neurons within connected populations are randomly associated according to specific connection probabilities (see Table 3). Each neuron receives inputs from within the network and from other brain regions (mainly the cortex) which have not been explicitly included in our model. These external inputs have been modeled as independent poissonian trains of pulses. The mean rates *ν*_*ext*_ of these signals have been adjusted for each population in order to ensure realistic population firing rates: FSN [10–20] Hz ([52], [48]), D1 and D2 (MSN) [0.05-2.5] Hz ([53]), GPe-TI [40-60] Hz ([38]), GPe-TA [5-15] Hz ([38]) and STN [12-20] Hz ([24]).

Note that the GPi and the SNr nuclei were not included in the simulated model since they do not present direct feedback to the other populations in the basal ganglia and hence cannot contribute to the generation of oscillations.

### 1.2 Neuron models

All neurons in the adopted network are adaptive, conductance based, point neurons ([54–57]). The STN, GPe-TI and GPe-TA populations are modelled as adaptive exponential neurons (`aeif_cond_exp` model in the code implementation) [58] and the dynamics of their membrane potential is governed by:

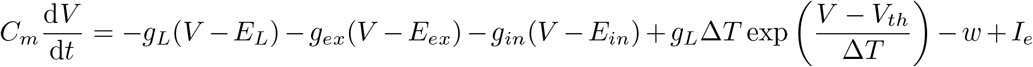

The striatal populations are modelled as adaptive quadratic neurons (`aqif_cond_exp` or `aqif2_cond_exp` model) [59] and their membrane potential evolves according to:

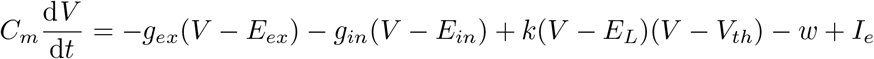

In both cases excitatory and inhibitory conductances present exponential decays:

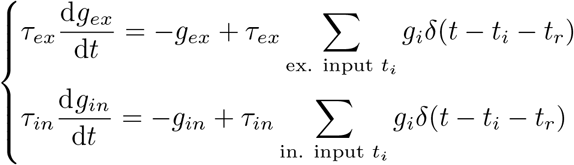

The evolution of the adaptation variable *w* is governed by ([60, 61]):

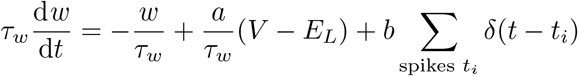

in the `aeif_cond_exp` and `aqif_cond_exp` model, while by:

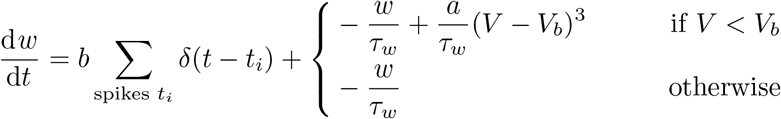

in the case of the `aqif2 cond exp` model.

To introduce variations in each neural population, the synaptic weight of the external poissonian input is not equal in all neurons. Rather, it is assumed to be uniformly distributed around its central value with an amplitude equal to the `dev-ext-weight` parameter. The adopted values of the parameters of each neuronal population are summarized in Table 2. The connectivity properties (delays, connection probabilities and synaptic weights) are reported in Table 3. These values have been adapted from the work of Lindhal and Kotaleski [51], here simplified neglecting synaptic plasticity, spatial restrictions on the connected populations and dopamine effects different from the ones described in section 1.5.

Note that the results of the model are not critically dependent on the exact values of the network parameters.

In the following, we will refer to the described network of the basal ganglia as the Complete Model.

### 1.3 Selection of relevant *β* oscillators

In order to identify the more relevant structures generating *β* oscillations, the following strategy has been pursued. Starting from the Complete Model (Fig 1), the relevance of each connection in generating oscillations in the *β* regime has been quantified referring to the decrease in the mean intensity of *β* oscillations following its elimination. In particular, for each connection *Source* → *Target* (*S* → *T*) in the adopted model, a simulation has been performed with the following properties:

- an *auxiliary* subnetwork *S*\* has been introduced with the same neuron model and size of *S* but with no input sources other than the external one;
- the parameter *I*_*e*_ of the *S*\* subnetwork has been adjusted so that the mean spiking-rate of *S*\* neurons was close to the one of the *S* population;
- the probability of connection between *S* and *T* neurons is set to 0;
- the *S*→ *T* connections are replaced by *S*\*→ *T* connections with the same probability and synaptic weight.

The relevance of the *S* → *T* connection has been thereby quantified by means of the ratio *R*(*S* → *T*) between the mean PSD with or without the (*S* → *T*) connection:

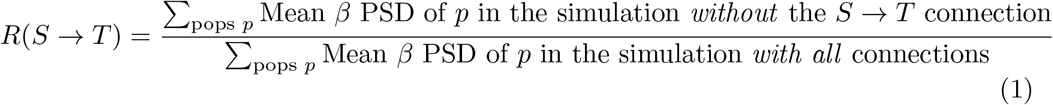

The connections whose ratio was lower were considered as more relevant in the generation of *β* oscillations. The major oscillators in the *β* regime have been identified on the basis of the selected connections. Self-inhibition connections have not been considered in this analysis since they generate oscillations at much higher frequencies [62].

### 1.4 Analysis of the effects of the coupling between *β* oscillators

As from our analysis emerged that two oscillators are mainly responsible for the onset of the pathological activity, we developed an *ad hoc* network to analyze the properties and the consequences of their interaction (see Fig 4A).

The GPe-TI nucleus is present in both oscillators, hence its neurons have been equally divided in two populations (GPTI-(A) and GPTI-(B)) belonging to the two different loops: neurons in GPTI-A are reciprocally connected to STN, while those in GPTI-B receive inputs from D2 and send outputs to FSN. These connections, which are represented as thick continuous lines in Fig 4A, present connection probabilities that are equal to the ones in the Complete Model.

In contrast, in order to modulate the coupling between the two oscillators, *inter*-loop connections (thin continuous line in Fig 4A) present connection probabilities which are modulated by a coupling parameter *ε*. Particularly, for each pair (*S,T*), the connection probability of each *S* neuron with each *T* neuron is set equal to:

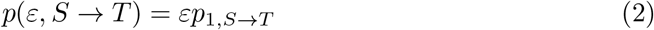

where *p*_1,*S*→*T*_ is the probability characterizing the corresponding connection in the Complete Model (see Table 3).

For every connection, synaptic weights are the same as in the Complete Model (and are not affected by *ε*). As a result, for *ε* = 0 the two loops are completely independent, while for *ε* = 1 the network presents realistic connectivity properties. Further, in order to ensure that the overall inputs to each nucleus remain the same when *ε* is changed, the *auxiliary* populations GPTI*, D2* e STN* have been introduced. Neurons in these nuclei receive constant and independent external input such that their mean spiking rates are equal to the mean spiking rates of the corresponding non-*auxiliary* populations. The *auxiliary* connections are shown as dashed lines in Fig 4A. While their synaptic weights are the same as in the original network, the connection probabilities *S*^*^ → *T* are set equal to:

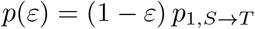

Hence, for *ε*=1 the auxiliary nuclei do not play any role; otherwise, they become external drives meant to compensate for the input that is not received from the *real* presynaptic populations. The neuron models and the parameters of the *auxiliary* populations are the same of the corresponding *real* populations (Table 2), except for *ν*_*ext*_ and the size *N* which is halved. As a consequence, the connection probabilities D2* →GPTI-A and STN*→ GPTI-B are doubled.

Since most nuclei in this simplified network receive fewer inhibition in comparison to the complete model, some adaptations in the external current *I*_*e*_ and the external poissonian input *ν*_*ext*_ have been introduced in order to keep the mean activity of the populations in the correct regime. Further, since all the non-*β* oscillators have been eliminated, *β* activity with the original connection weights is dominant for all *ε*; to study the process of synchronization in these states, it has been necessary to modulate the intensities of the connections constituting the two loops.

The complete list of the introduced adaptations are listed in Table S1. In the following, and in contrast to the original Complete Model (see Section 1.1), the network obtained after the listed assumptions and simplifications will be labelled Simplified Model.

### 1.5 Dopamine depletion modeling

In order to investigate the role of dopamine and accounting for its effects on the mean spiking rate of neurons into striatal populations ([10]), the severity of dopamine depletion has been modelled in the Simplified Network by the modulation of the intensity of the external input rate towards the D2 population. More precisely, the external input rate for this population (and the corresponding auxiliary population D2*) has been modulated according to:

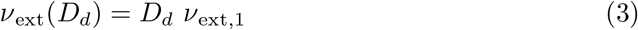

where *ν*_ext,1_ is the reference value of the input rate (see Table 2) and *D*_*d*_ is the parameter regulating the severity of the condition of dopamine depletion (the higher is *D*_*d*_, the more severe is the condition). Throughout the analysis, *D*_*d*_ varies in an interval such that the D2 activity ranges from *∼*0.05 to *∼*4 Hz. The effects of dopamine depletion will be also studied in the Complete Model applying the same modulation.

### 1.6 Spectral Analysis

We measured the activity of each population as the time series of its firing rate computed over time bins of one millisecond. For each nucleus we performed a constant detrend and computed the Power Spectral Density (PSD) of the activity. In order to reduce the effects of noise, the Welch method has been applied (we employed the `signal.spectrogram` function from `scipy` with `N_parseg`= 2000, `N_overlap`= 1000 and Tukey window: `alpha`=0.25); further, for each case studied, 4 simulations have been performed in order to estimate the standard deviation of each computed quantity.

Starting from the PSD of the activity of each population *i* within the network, we defined:

- its mean frequency of oscillation:

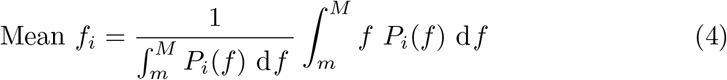

- its mean value in the *β* regime:

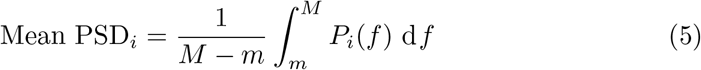

where *P*_*i*_(*f*) is the PSD of the nucleus activity. The values of *m* and *M* have been fixed to 8 and 24 Hz respectively, within the *β* regime and coherently with the natural frequencies of the two oscillators.

The Mean PSD_*i*_ is a measure of the intensity of *β* oscillations within each considered nuclei. We highlight that this measure is *biased* : in the presence of constant activity (no *β* oscillations), fluctuations around the mean are present and the related spectrum is affected by noise. In order to eliminate this bias we considered the corrected quantity:

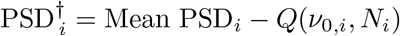

where *Q*(*ν*_0,*i*_, *N*_*i*_) is the Mean PSD of the activity of a population of *N*_*i*_ neurons with constant mean activity *ν*(*t*) = *ν*_0,*i*_. The value of *Q*(*ν*_0,*i*_, *N*_*i*_) has been estimated computing the PSD of a fictitious activity in which the number 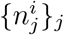 of spiking neurons in each bin is given by independent extractions from a Bernoulli distribution:

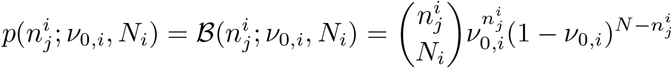

### 1.7 Analysis of the limit of large population

In order to verify whether the properties that we obtained persist in the limit of large populations (the realistic size of each population is approximately 10^3^ times the adopted sizes [63, 64]) we repeated all the analysis with each population size multiplied by a factor *n ∈* [1, 2, 4]:

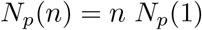

where *p ∈* [FSN, D1, D2, GPe-TI/TA, GPTI-A/B, STN, D2*, STN*, GPTI*] are the different populations and *N*_*p*_(1) is the size of population *p* listed in Table 1. In order to have comparable results for different values of *n*, the PSDs have been computed over *N*_*p*_(1) neurons also in the case of *n* ≠ 1. Further, in case of *n* ≠ 1, all connection probabilities were scaled in such a way that the mean number of presynaptic neurons per target neuron was constant for each connection.

**Table 1.** Reference size and neuron model of each population. Note that size was modulated in a subset of the analysis.

| SubNetwork | $N$ | model |
| --- | --- | --- |
| D1 (MSN) | 6000 | aqif_cond_exp |
| D2 (MSN) | 6000 | aqif_cond_exp |
| FSN | 420 | aqif2_cond_exp |
| GPe-TA | 264 | aeif_cond_exp |
| GPe-TI | 780 | aeif_cond_exp |
| STN | 408 | aeif_cond_exp |

**Table 2.**
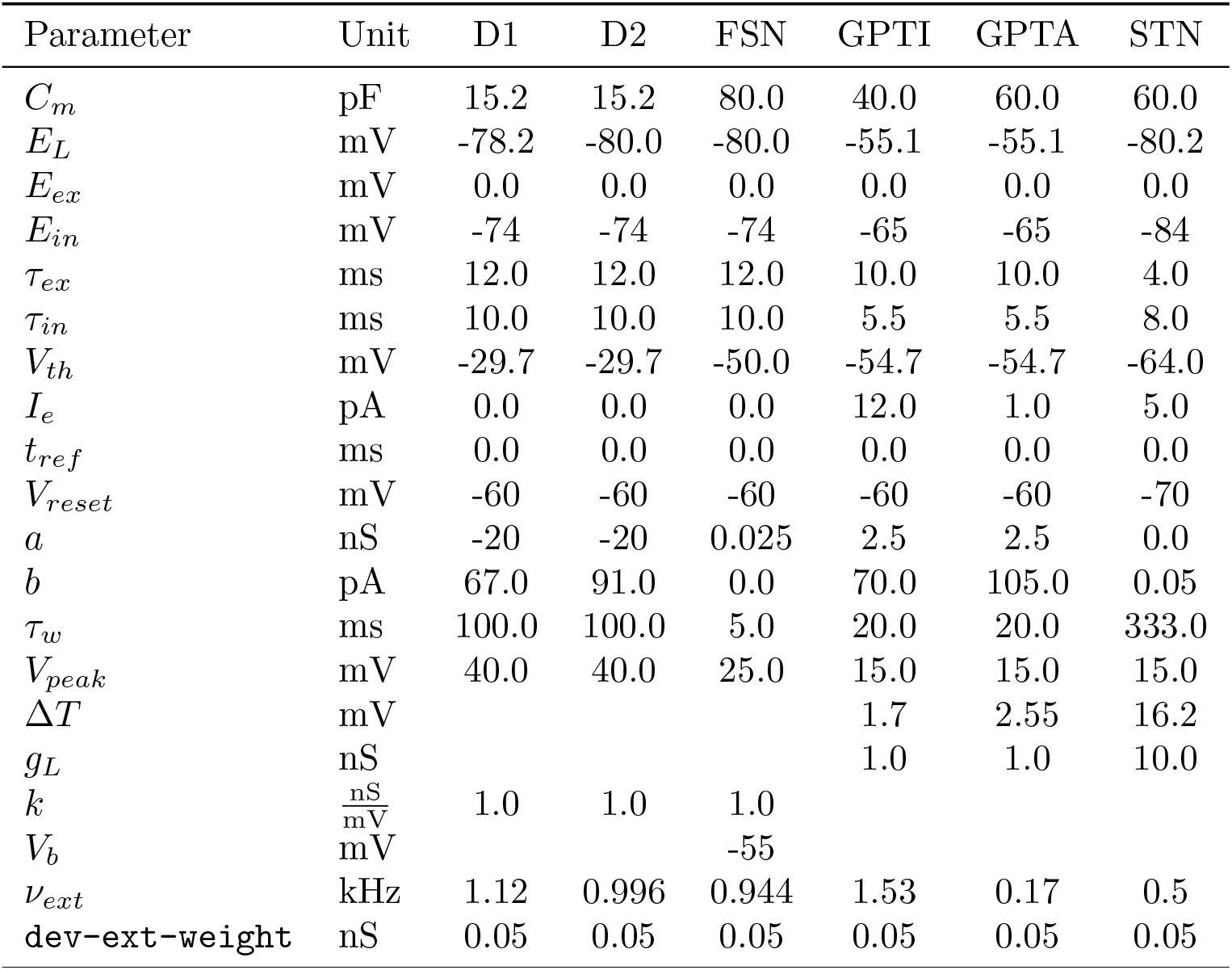
Adopted values of the parameters of each neuronal population.

**Table 3.** Connectivity properties of the adopted model. Type I represents inhibitory connections while type E represents excitatory connections.

| Source | Target | Probability | Delay [ms] | Type | synaptic weight [nS] |
| --- | --- | --- | --- | --- | --- |
| D1 | D1 | 0.0607 | 1.7 | I | 0.12 |
|  | D2 | 0.0140 | 1.7 | I | 0.30 |
| D2 | D1 | 0.0653 | 1.7 | I | 0.36 |
|  | D2 | 0.0840 | 1.7 | I | 0.20 |
|  | GPTI | 0.0833 | 7. | I | 1.28 |
| FSN | D1 | 0.0381 | 1.7 | I | 6.60 |
|  | FSN | 0.0238 | 1.0 | I | 0.50 |
|  | D2 | 0.0262 | 1.7 | I | 4.80 |
| GPTI | GPTI | 0.0321 | 1.0 | I | 1.20 |
|  | GPTA | 0.0321 | 1.0 | I | 0.35 |
|  | FSN | 0.0128 | 7.0 | I | 1.60 |
|  | STN | 0.0385 | 1.0 | I | 0.08 |
| GPTA | D1 | 0.0379 | 7.0 | I | 0.35 |
|  | D2 | 0.0379 | 7.0 | I | 0.61 |
|  | FSN | 0.0379 | 7.0 | I | 1.85 |
|  | GPTA | 0.0189 | 1.0 | I | 0.35 |
|  | GPTI | 0.0189 | 1.0 | I | 1.20 |
| STN | GPTA | 0.0735 | 2.0 | E | 0.13 |
|  | GPTI | 0.0735 | 2.0 | E | 0.42 |
| ext | D1 | 1.0 | 0.0 | E | 0.45 |
|  | D2 | 1.0 | 0.0 | E | 0.45 |
|  | FSN | 1.0 | 0.0 | E | 0.50 |
|  | GPTI | 1.0 | 0.0 | E | 0.25 |
|  | GPTA | 1.0 | 0.0 | E | 0.15 |
|  | STN | 1.0 | 0.0 | E | 0.25 |

### 1.8 Numerical methods

The code for the simulations has been developed in C++. In combination with this code, a `python3` module has been implemented in order to import and preliminarily analyse the results of the simulations. The implemented code is available on [GitLab](https://gitlab.com/andrea.ortone/basal_ganglia_model) and the documentation is published on readthedocs.org. The numerical integration of the equations describing the evolution of each neuron in the simulated network are performed with a fixed time step *h* = 0.1 ms and applying the 4^*th*^ order Runge-Kutta procedure ([65]). In order to generate random numbers the [PCG library](https://www.pcg-random.org/index.html) ([66]) has been employed. The duration of each simulation has been fixed to 10 000 ms plus a warm-up interval of 500 ms which has been eliminated from the analysis.

## Results

With the aim of untangling the dynamics leading to the onset of pathological *β* oscillations, we first identified the main subnetworks generating *β* resonances, and then studied the effects of their interactions. In order to highlight the role of the pathology, we finally investigated the role of dopamine depletion in shaping the spectral dynamics of the system.

### 2.1 Identification and characterization of *β* oscillators

In order to identify the connections mostly contributing to the generation of *β* oscillations, we considered the Complete Model of the basal ganglia (see Fig 1 and Methods section 1.1) and computed the residual *β* power *R*(*c*) associated to the removal of each connection *c* from the network (see 1.3 for details). According to the definition in Eq. (1), the more relevant connections are expected to be associated to lower values of the ratio *R*, hence we fixed a threshold 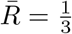 and focused on the connections with 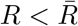. As a result, five connections were identified as main responsible for the onset of *β* activity (Fig 2A). These connections set up two distinct oscillators, one involving FSN, D2 and GPe-TI nuclei (STR loop, Fig 2B) and the other involving STN and GPe-TI nuclei (STN loop, Fig 2C). The spectral analyses of the activity of the two oscillators (isolated from the other nuclei by setting *ε* = 0, see Methods) showed that:

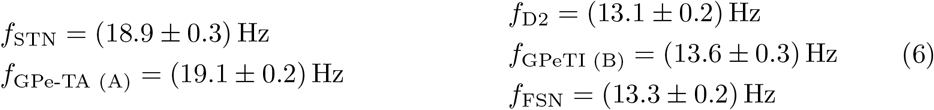

hence the STR loop has a proper frequency *f*_STR_ *∼* 13.3 Hz while STN naturally oscillates at *f*_STN_ *∼*19 Hz.

**Fig 2.**
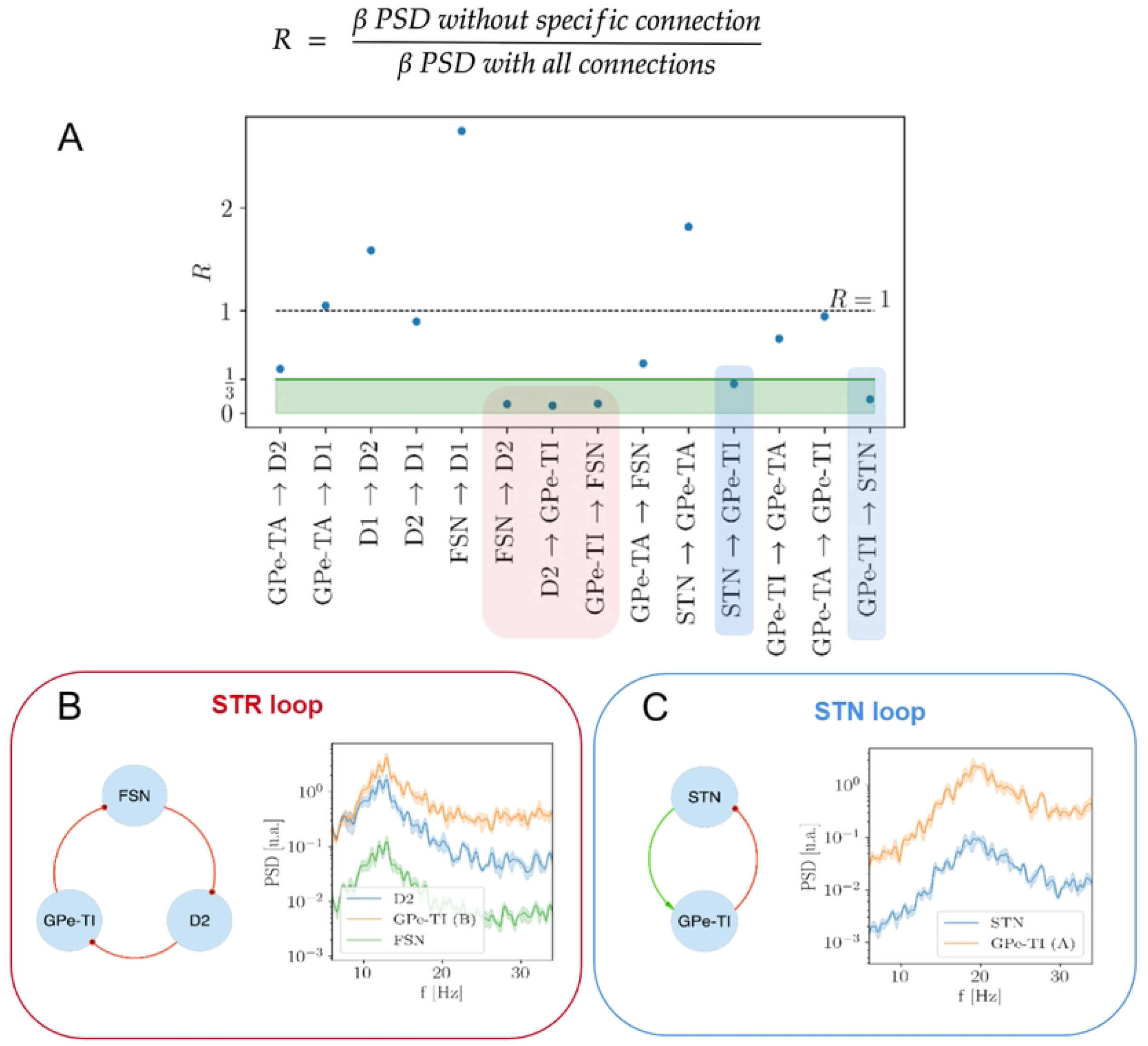
Identification of most relevant oscillators and investigation of their spectral properties. **(A)** Rationale about the selection of salient nodes in the basal ganglia network for a single simulation run. The nodes, taken together, define the STR and the STN loops (in blue and red shadow respectively). **(B)** STR loop with FSN, GPe-TI and D2 nuclei, connected with feedforward inhibitory projections (in red) and with their natural mode. **(C)** STN loop with STN and GPe-TI nuclei, connected with excitatory feedforward (in green) and inhibitory feedback projections (in red) and their natural mode.

Since both loops contain the GPe-TI population, in the Complete Model of the basal ganglia the two loops interact with one another. The consequences of this interplay will be the objective of the next sections.

### 2.2 Loops interplay

The results highlighted in the previous section suggest that the main responsible for the onset of *β* activity in the adopted model of the basal ganglia are the two oscillators set up by the STR and STN loops. In order to deepen the properties of the interaction between these oscillators, we now consider the Simplified Model (Fig 4A). In this section, we particularly focus on the consequences of increasing the value of the coupling *ε* which regulate the intensity of the connections between the two oscillators (see Methods section 1.4).

Interplay variability resulted in fundamental changes in the spectral dynamics of the network (Fig 3): for very low values of *ε* the Str and STN loops, measured through the D2 and STN nuclei respectively, exhibit a power spectrum which is not dissimilar from the case of the independent oscillators (*ε* = 0) (Figs 2B and 2C). As the value of the *ε* parameter is increased, the PSDs of both populations become affected by the presence of the other resonance and present a broader peak which increasingly include the intermediate region between the natural frequencies of the two independent oscillators. Finally, for high values of *ε ∼*0.85, the oscillators complete their process of synchronization: the two separate resonances disappear and a single prominent peak emerges at an intermediate frequency.

**Fig 3.**
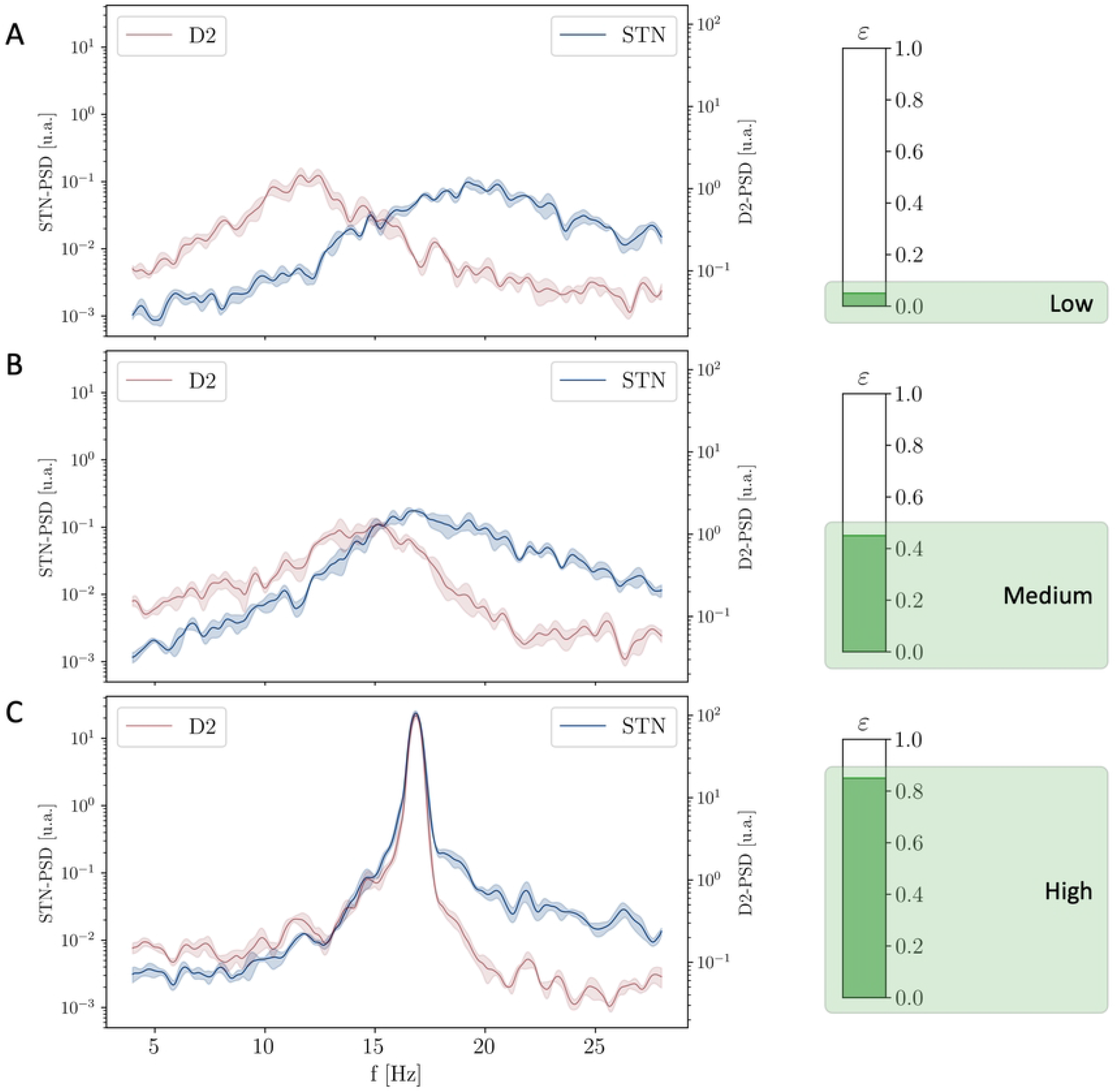
Evolution of the PSD of the STN and D2 populations for increasing values of the coupling strength *ε*. **(A)** PSD of STN and D2 nuclei related to a low level of coupling (*ε ∼* 0.05) showing asynchronous states. **(B)** PSD of both STN and D2 nuclei in relation with an intermediate level of coupling (*ε ∼* 0.45) showing larger peaks increasingly including the intermediate region. **(C)** PSD of STN and D2 nuclei related to a high level of coupling (*ε ∼* 0.85) showing the emergence of a common oscillatory mode.

In order to highlight this behaviour, we computed the average frequency of the STR and STN loops (with specific measurements respectively at the D2 and STN nuclei) as a function of *ε* (see Methods section 1.6). As expected, the two frequencies start from the natural frequencies of the two oscillators (13 *∼* Hz and 19 *∼*Hz in the region of *ε* = 0) and, as the interplay between the two loops increases, converges to an intermediate common frequency *f* 16 *∼* Hz (*ε* = 1, see Fig 4B). Even more interestingly, the process of synchronization is associated to a strong increase in the intensity of the activity in the *β* regime (Fig 4C).

**Fig 4.**
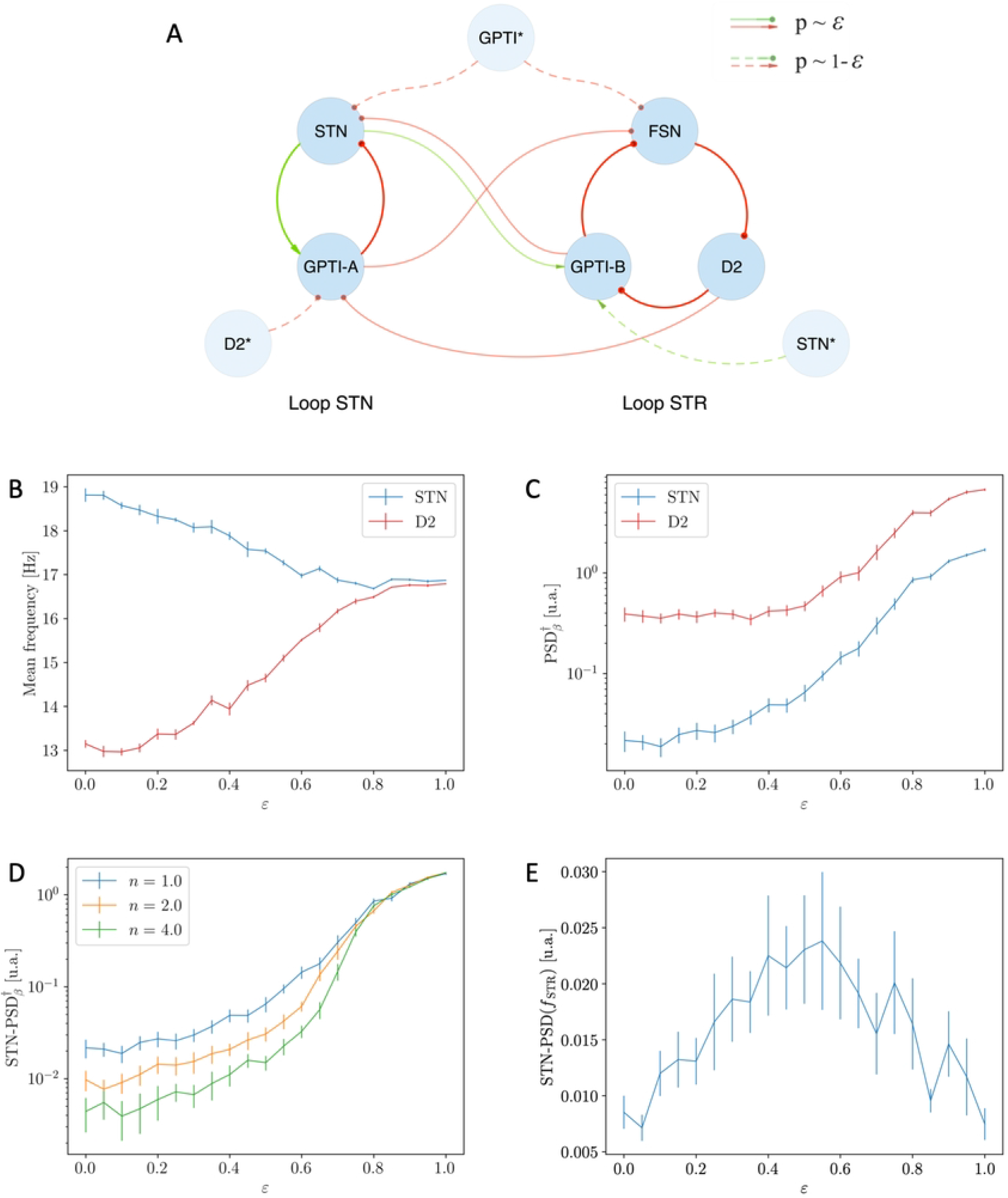
Effects of the coupling strength *ε* on the Simplified Model. **(A)** Schematic representation of the Simplified Model: the modulation of the coupling strength between the STR and STN loops is obtained by varying the connection probabilities *p*(*ε, S*→ *T*) = *εp*_1,*S*→*T*_ (see Eq. (2)) of *inter*-loops connections. **(B)** Mean frequencies of STN (blue) and D2 (red) nuclei as function of the coupling strength *ε*: the two mean frequencies converge due to the increase of the coupling strength between the two oscillators. **(C)** Unbiased measure of the intensity of *β* activity as a function of *ε*: note the remarkable growth of the intensity in the synchronous regime. **(D)** Unbiased measure of the intensity of the STN *β* activity as a function of *ε* for different values of the population size *n* (blue n=1.0, orange n=2.0, green n=4.0): the intensity of *β* activity is preserved only in the synchronous regime. **(E)** Value of the STN-PSD nuclei computed at the natural frequency of the STR oscillator as a function of the coupling strength. These results are associated with the analogous in panels B and C of Fig S2.

In order to verify whether the properties highlighted above persist in the limit of large populations, we repeated the analysis with the size of each population multiplied by a factor *n ∈* [2, 4] (see Methods sec 1.7). The results of this analysis highlight two different regimes (Fig 4D):

- in case of low synchronization (*ε ∼* 0) the intensity of *β* oscillations goes to zero in the large *n* limit;
- in case of high synchronization (*ε ∼* 1) the intensity of *β* activity is preserved.

On the one hand, these results confirm that the establishment of a state of complete synchronization and the consequent onset of prominent *β* activity in the model are stable for variations in populations size; on the other hand, they suggest that single loop models are not capable of explaining strong *β* activity in the limit of large populations. A further insight on the process of synchronization is finally captured by the value of PSD_STN_(*f*_STR_) (i.e., the power spectral density of the STN population computed at the natural frequency of the STR loop; Fig 4E). Particularly, for low coupling, this value is low as STN uniquely oscillates at the natural frequency of the STN loop (Fig 3A). In the intermediate regime (*ε ∼* 0.5) higher values are registered since the activity of the STN nucleus becomes more affected by the STR oscillator (Fig 3B). Finally, for *ε ∼* 1, the value of PSD_STN_(*f*_STR_) decreases again as the two oscillators synchronize to a novel frequency which is different from both the natural ones (Fig 3C).

### 2.3 Role of dopamine in *β* synchronization dynamics

So far we described the effects of synchronization as a result of increasing the value of the coupling *ε* between the two *β* oscillators. In the real network, however, there are no certain evidences of the fact that the intensity of the connections between the loops significantly increases due to the emergence of Parkinson’s Disease (i.e. the value of *ε* is fixed). This fact implies that there must be something else, within the network, which is responsible for the increase of the interaction between the two oscillators and hence drives the system towards higher degrees of synchronization.

In this section, we focus on the effects of increasing the condition of dopamine depletion by regulating the value of the *D*_*d*_ parameter (see Methods section 1.5); in all the following studies the value of *ε* will be fixed.

#### The role of dopamine in the case of uncoupled oscillators

As a preliminary study, we first consider the effects of *D*_*d*_ in the Simplified Model and fixing the value of *ε* = 0.00.

In this case, the main effect of increasing the value of *D*_*d*_ (i.e. the severity of the condition of Dopamine Depletion, see Methods and Eq. (3)), is that of enhancing the intensity of the oscillator set up by the STR loop: in the healthy status (*D*_*d*_ *∼* 0.9) the striatum resonance is almost absent; in contrast, as *D*_*d*_ is increased, the intensity of the oscillations strongly increases (see Figs 5B and 5D). Note also that, since the two oscillators do not interact, the system does not undergo any process of synchronization: the mean frequency of the STN nucleus sticks close to *∼* 18 Hz, while that of the D2 nucleus moves from *∼* 16 Hz, that is mean value between 8 and 24 (see Methods and Eq. (4)), to the natural frequency of the D2 oscillator at *∼* 13 Hz (see Fig 5C).

**Fig 5.**
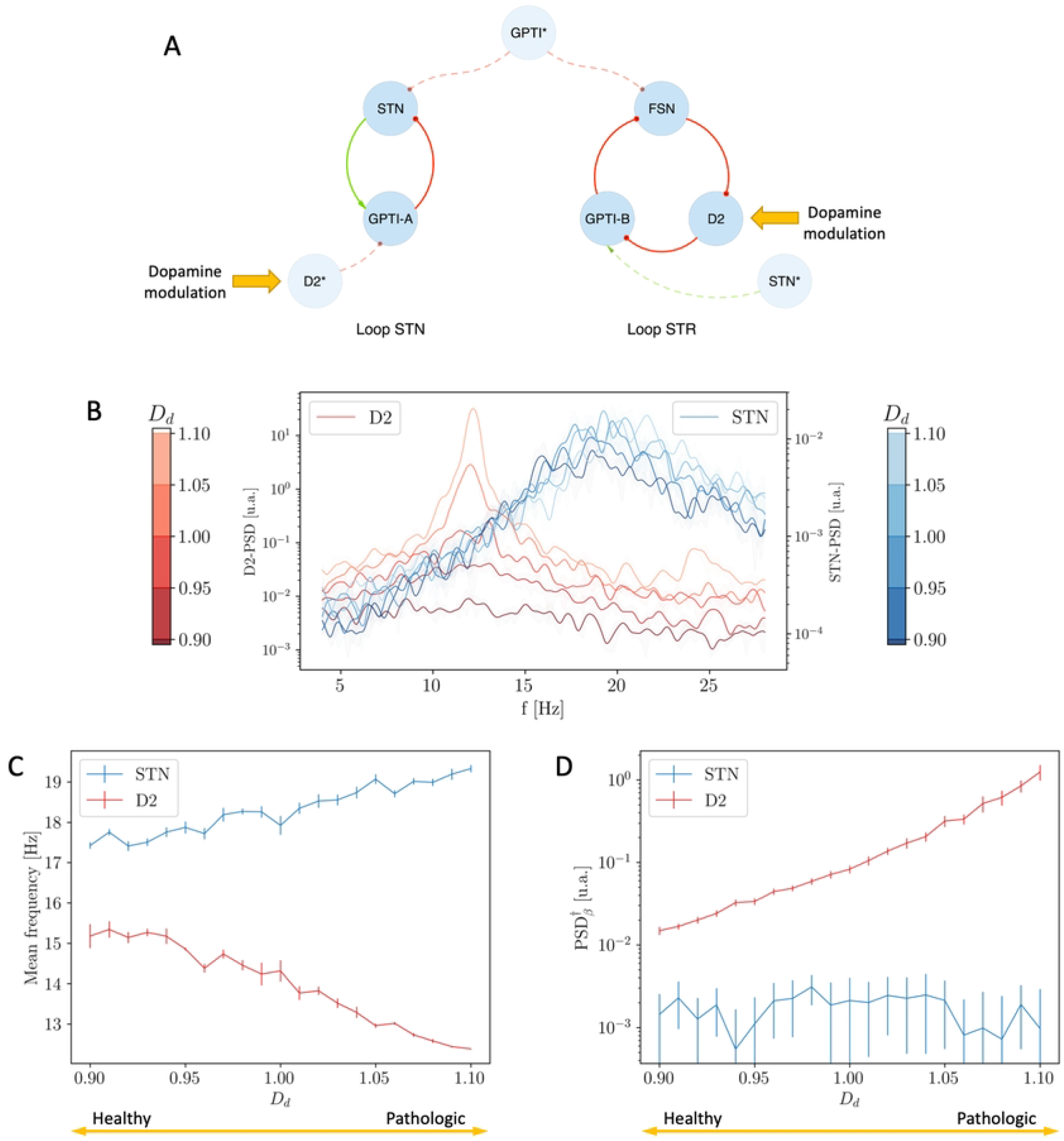
Effects of Dopamine Depletion *D*_*d*_ on the Simplified Model (*ε* = 0.00). **(A)** Schematic representation of the Simplified Model in the non-interacting case: note the highlighted targets of Dopamine modulation. **(B)** Evolution of the PSDs for increasing values of *D*_*d*_ and for the D2 (red) and STN (blue) populations. **(C)** Mean frequencies of STN (blue) and D2 (red) nuclei as function of dopamine modulation *D*_*d*_: the two mean frequencies do NOT converge due to the increase of *D*_*d*_. **(C)** Unbiased measure of the intensity of *β* activity as a function of *D*_*d*_: note that the intensity of the D2 resonance grows for increasing values of *D*_*d*_.

#### The role of dopamine in the Simplified Model

We continue considering the Simplified Model but we now introduce the coupling between the two oscillators by setting the value of *ε* = 0.75.

In contrast to the previous case, the increase in the intensity of the STR Loop leads the two oscillators to undergo a process of synchronization: for low values of *D*_*d*_ the two oscillators interact poorly (the STR oscillator is almost absent and only the STN resonance at *∼* 18 Hz characterizes the status of the system; see Fig 6A); as the value of *D*_*d*_ is increased, the enhancement in the intensity of the STR oscillator implies higher degrees of interaction between the two loops and consequently leads to the emergence of a single resonance at an intermediate frequency (*∼* 16 Hz; see Figs 6B and 6C).

**Fig 6.**
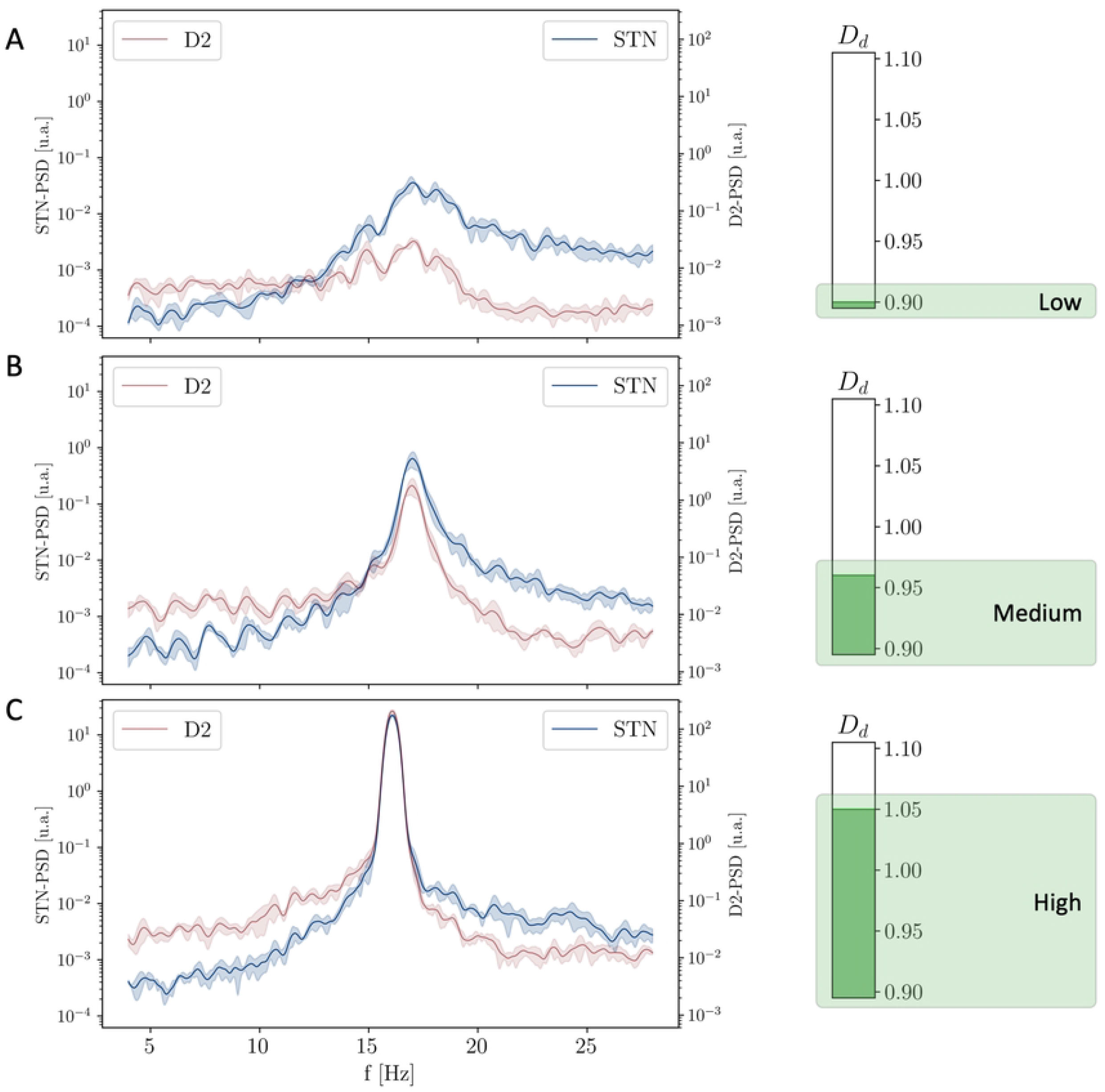
Evolution of the PSD of the STN and D2 populations for increasing values of Dopamine Depletion *D*_*d*_. PSD of both STN and D2 nuclei related to low **(A)**, intermediate **(B)** and high **(C)** levels of dopamine depletion (*D*_*d*_=0.9,0.97 and 1.05 respectively): for low values of *D*_*d*_ the system is characterized by the only STN-loop resonance; the increase of *D*_*d*_ leads to higher degrees of interaction and to the emergence of a unique resonance at an intermediate frequency.

Similarly to the increase of *ε* (section 2.2), the increase of *D*_*d*_ determines the convergence of the mean frequencies of oscillation of the STN and D2 populations (Fig 7B) and a remarkable increase in the intensity of the pathological oscillations (Fig 7C).

**Fig 7.**
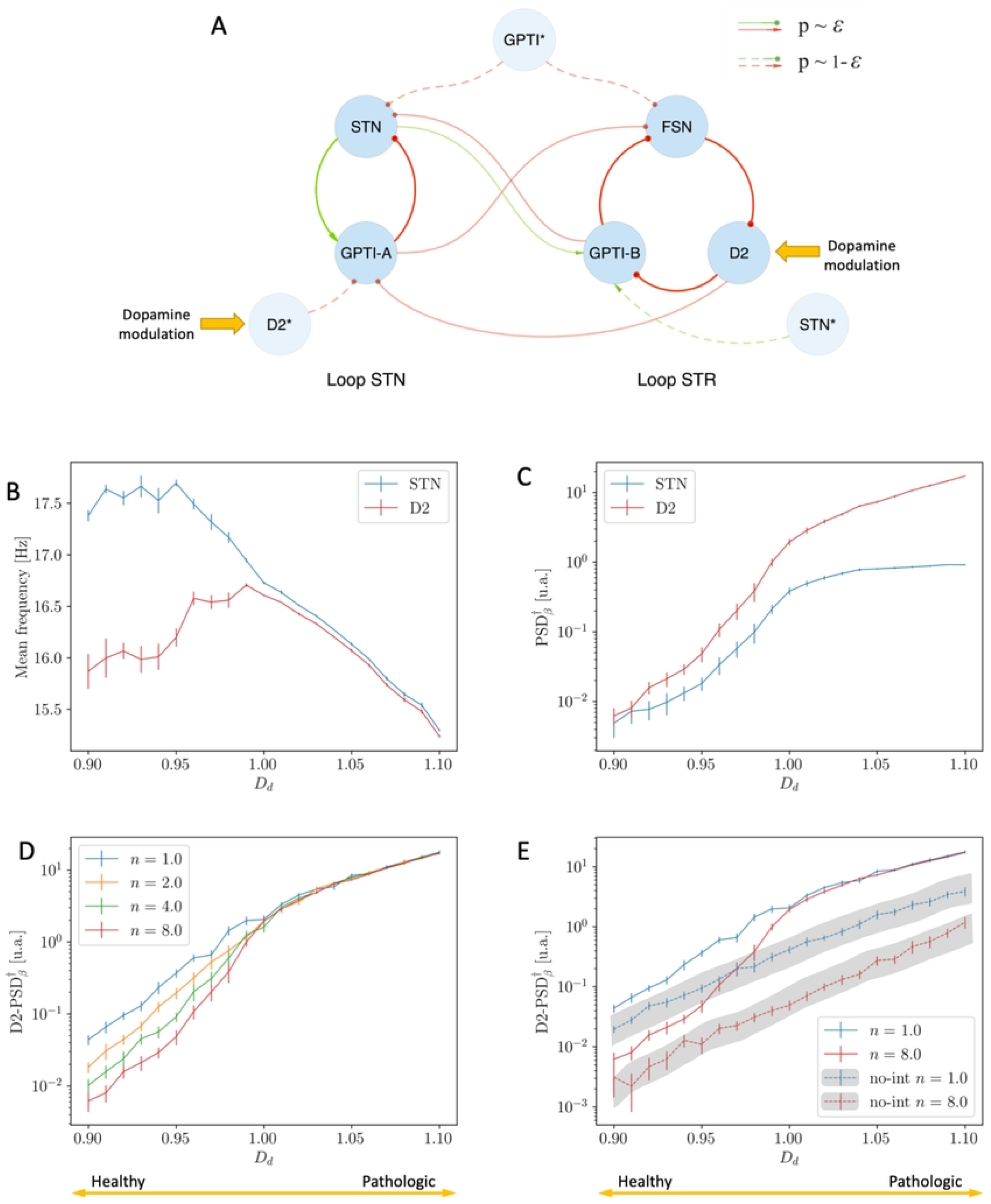
Effects of Dopamine Depletion *D*_*d*_ on the Simplified Model (*ε* = 0.75). **(A)** Schematic representation of the Simplified model in the interacting case: note the highlighted targets of Dopamine modulation. **(B)** Mean frequencies of STN (blue) and D2 (red) nuclei as function of dopamine modulation *D*_*d*_: the two mean frequencies converge due to the increase of *D*_*d*_. **(C)** Unbiased measure of the intensity of *β* activity as a function of *D*_*d*_: note the remarkable growth of the intensity in the synchronous regime. **(D)** Unbiased measure of the intensity of the D2 *β* activity as a function of *D*_*d*_ for different values of the population size *n* (blue *n*=1.0, orange *n*=2.0, green *n*=4.0, blue *n*=8): the intensity of *β* activity is preserved only in the synchronous regime. **(E)** Unbiased measure of the intensity of the D2 *β* activity as a function of *D*_*d*_ for *n* = 1 (blue) and *n* = 8 (red) and comparison between the interacting (*ε* = 0.75, continuous lines) and non-interacting condition (*ε* = 0.00, dashed lines): the intensity of *β* activity is preserved if and only if the two oscillators are synchronized. These results are associated with the analogous in panels B and C of Fig S3.

Further, the analysis of the limit of large populations shows that:

- if *ε* = 0.75 and *D*_*d*_ *∼* 1.1 (synchronized oscillators), the intensity of *β* activity is preserved (Fig 7D, right);
- if *ε* = 0.75 and *D*_*d*_ *∼* 0.9 (non-synchronized oscillators), the intensity of *β* activity decreases (Fig 7D, left);
- if *ε* = 0.00, for any value of *D*_*d*_ (non-synchronized oscillators), the intensity of *β* activity decreases (Fig 7E).

In accordance with previous results, this proves that the intensity of *β* activity in the large *n* limit is only preserved when the two oscillators are in the synchronous state (*ε* = 0.75 and *D*_*d*_ *∼* 1.1).

Beside that, the comparison between the uncoupled (*ε* = 0.00) and coupled (*ε* = 0.75) cases highlights that the increase in the intensity of *β* activity is stronger in the latter case (Fig 7E).

The results proposed in this section concerning the role of dopamine in the case of the Simplified Model showed interesting changes in the functional and spectral dynamics of the two loops. The greater the presence of dopamine (healthy condition), the lower the state of synchronisation between the STR and STN loops, and the greater their functioning around their natural resonance regime. Conversely, the lesser the presence of dopamine (pathological condition), the greater the state of synchronisation of the two loops, and the more the activity of the network is characterized by a single, prominent oscillation mode in the *β* regime.

#### Role of dopamine in the Complete Model

In all previous analyses we focused on the properties of the emergent *β* activity in the context of the Simplified Model. In that case, we showed that the effects of dopamine depletion and the consequent process of synchronization of the two oscillators set up by the STR and STN loops play a major role in the onset of pathological, prominent *β* activity. In line with those results, in this section we will show that all the highlighted properties are preserved in the Complete Model of the basal ganglia (see Fig 8A). Particularly, even in this model, as the value of *D*_*d*_ is increased, the mean frequencies of the D2 and STN populations converge (Fig 8B) and the intensity of *β* oscillations increases and presents the characteristic speed up due to completion of the synchronization process (Fig 8C).

**Fig 8.**
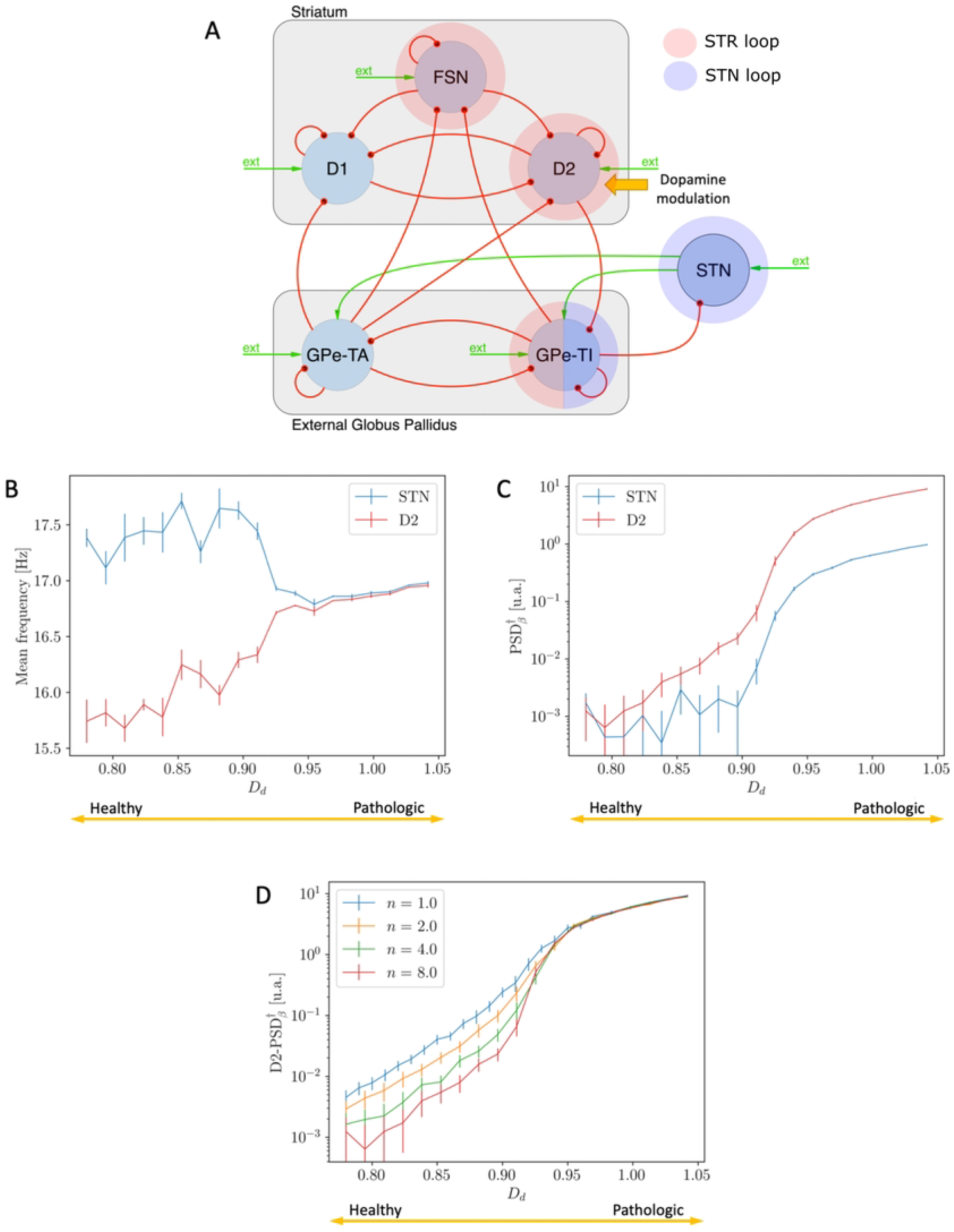
Effects of Dopamine Depletion *D*_*d*_ on the Complete Model. **(A)** Schematic representation of the Complete Model: note the highlighted target of Dopamine modulation. **(B)** Mean frequencies of STN (blue) and D2 (red) nuclei as function of dopamine modulation *D*_*d*_: the two mean frequencies converge due to the increase of *D*_*d*_. **(C)** Unbiased measure of the intensity of *β* activity as a function of *D*_*d*_: note the remarkable growth of the intensity in the synchronous regime. **(D)** Unbiased measure of the intensity of the D2 *β* activity as a function of *D*_*d*_ for different values of the population size *n* (blue *n*=1.0, orange *n*=2.0, green *n*=4.0, red *n*=8): the intensity of *β* activity is preserved only in the synchronous regime. Analogous results about STN *β* activity are associated with Panel B of Fig S4.

Finally, and in accordance with previous results, the size analysis confirms that our model is capable of capturing prominent *β* activity also in the limit of large populations (Fig 8D).

Please note that in all the studies regarding the effects of dopamine depletion (Figs 5, 6, 7B, 7C, 8B and 8C) we showed the results related to the case *n* = 8. This has been necessary to capture the relevant properties of the system in a clearer way (for example, the reader can compare the speed up in the increase of *β* activity in case of *n* = 1 and *n* = 8 shown in Fig 7E). This choice does not affect the results of the large *n* limit analysis.

#### *β* bursting activity

The presented results have been obtained by mediating over different time intervals and different simulations (see Methods section 1.6). Performing this operation denied us the possibility of investigating the transient properties of the network.

One of the most important feature of the pathological *β* activity in the basal ganglia is that it presents a strongly phasic characterization ([19, 20]).

As shown in Fig 9, our model is capable of capturing this characterization. In particular, for intermediate values of *D*_*d*_ (*D*_*d*_=0.90), the system gains temporary access to the synchronous condition and the two oscillators continuously switch between the synchronous and asynchronous states. Moreover, when the value of Δ*f* (*t*) = *f*_STN_(*t*) − *f*_D2_(*t*) decreases (meaning that the system spends more time in the synchronous status), the intensity of the *β* oscillations increases (see the red-green and green-red matching of points related to the same time intervals in the two inferior subplots of Fig 9). This result is in accordance with the previous considerations and indicate once again that the more the two oscillators interact, the more the system spends time in the synchronous state and the higher is the intensity of *β* activity within the network.

**Fig 9.**
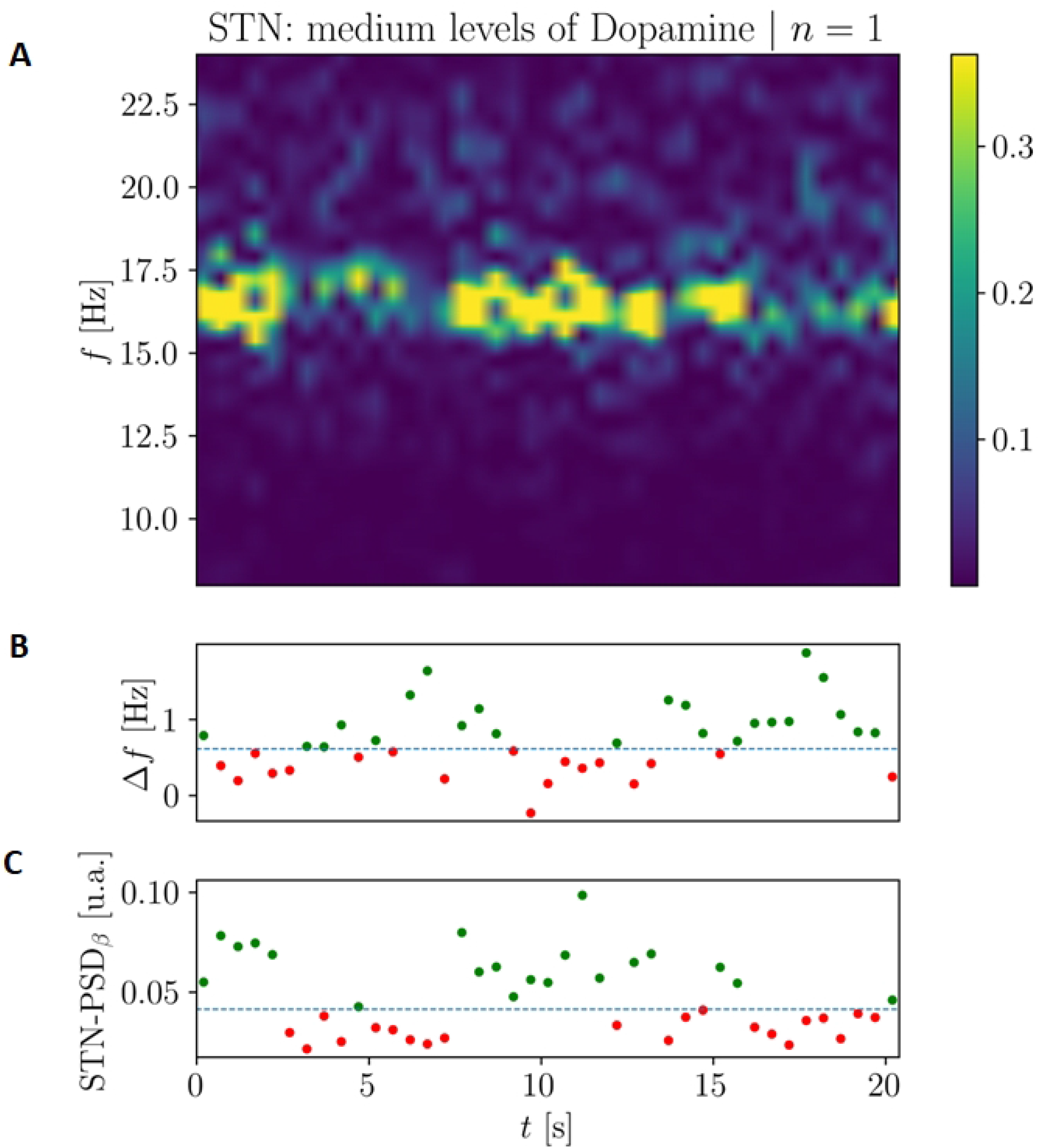
Bursting dynamics for intermediate values of dopamine (*D*_*d*_ = 0.90) in the case of the complete model of the basal ganglia. **(A)** Periodogram of the STN population activity highlighting the bursting characterization along a period of 20 seconds. **(B)** Difference between the instantaneous mean frequency of STN and D2 nuclei. **(C)** Instantaneous values of PSD for the STN nucleus. The dashed lines in the inferior subplots represent the median values of Δ*f* and *β* STN Mean PSD respectively.

## Discussion

We characterized the interplay between two loops generating *β* oscillations within the basal ganglia: one between GPe and striatum and one between GPe and STN. While for high values of dopamine the two loops are both weak and act largely as independent, when dopamine is depleted the striatopallidal loop increases its strength, and this leads to synchronization and mutual reinforcement between the two loops and eventually to strong oscillations in the whole basal ganglia network. We suggest that this mechanism plays a key role in the generation of abnormal *β* oscillations in Parkinson’s Disease.

Our findings are coherent with state-of-the-art experimental results, and shed light on the underlying mechanisms. D2 neurons are known to increase their firing activity and to strongly synchronize in the *β* range in dopamine-depleted conditions [13]. Our modeling work suggests a mechanism in which the former effect leads to the latter due to the internal resonances of the BG network. The relevance of GPe in the generation of *β* oscillations was recently proved by a work of Mallet group [42]. First, the cortical origin of *β* oscillations was ruled out by observing that optogenic cortical silencing did not affect *β* power. Interestingly, also STN activity suppression had a small effect on *β* power, suggesting that STN might only have, in the words of the authors “a supportive role” in the generation of abnormal *β* oscillations, which where instead suppressed by GPe silencing. These results are coherent with our view in which the main driver of abnormal *β* oscillations is the increasing power of the striatopallidal *β* loop induced by dopamine depletion effects on D2, although oscillations are enhanced by the synchronization with the subthalamopallidal loop. It is remarkable that in [42] this synchronization is investigated inducing artificial GPe *β* oscillations and observing that the STN loop synchronize to them. Moreover, our results are coherent with classic results highlighting the role of the STN-GPe loop in generating *β* oscillations [21] as this loop indeed exists and does play an important role. The key difference is that in our view this loop is not simply modulated by the intensity of the striatal inhibition to GPe, but rather by the intensity of the *β* oscillations of the striatopallidal loop.

Most previous computational models on the onset of *β* oscillations (e.g., [26, 34, 67]) described how the STN-GPe loop was originating the pathological oscillations but focused only on perturbations induced by modulations of the intensity from striatal population to pallidum [29], without taking into account the pallidostriatal feedback that leads to the striatopallidal *β* loop. In a relevant work, Corbit and colleagues introduced a detailed small scale model of the striatopallidal loop [50] and showed how it generated *β* oscillations. Later models, including [51] which was the model laying the ground for this work, implemented then this loop in their network descriptions ([68, 69]), but did not explicitly address how the two loops interacted in originating *β* oscillations, rather focusing on action selection. Interestingly, a recent study by the same group focusing on transient responses of the basal ganglia network [70] also captured *β* activity, but did not address the question of its origins.

In a key paper, Tachibana et Al [71] studied the effects of (locally) blocking different connections of the STN-GPe circuit on the intensity of *β* activity. As accurately described in [72], the strong constraints highlighted in [71] rejected most of the previous hypotheses on the origin of *β* oscillations (see Introduction). Differently from those cases, thanks to the presence of two loops and to the effects of their synchronization (which are not affected by local blocks of the connections), the results of our model are coherent with the evidences in [71]. Further, and differently from the mechanisms proposed by Pavlides et al [72], our model does not need to include cortical populations.

As a final remark, the presented size analyses highlight that the intensity of *β* activity is preserved when the two oscillators are in the synchronous regime. On the one hand this confirms that our model is cable of reproducing *β* activity also in the limit of large populations; on the other hand it suggests that single loop models ([29, 34, 50]) may not be capable of explaining prominent *β* activity.

In future works we will investigate how our understanding of the mechanisms underlying the pathological oscillatory behaviour in Parkinson’s disease can lead to improved therapies. For instance, the dopamine depletion values in our model could both increase due to the progression of PD and decrease due to the action of dopamine agonists, but to proper model this latter aspect we have to take into account several other pathophysiological aspects [73]. Moreover, it would be interesting to simulate the action of Deep Brain stimulation as in [67, 74–76] also in our network to evaluate the input of different targets on suppressing the *β* oscillations and compare it to experimental results [77–79].

## Acknowledgments

AM was supported by a Clinical Grant of the Marlene and Paolo Fresco Foundation for Parkinson’s Disease and by internal funds of Scuola Superiore Sant’Anna. AV was supported by the PREVIEW project (PRedicting the EVolution of SubjectIvE Cognitive Decline to Alzheimer’s Disease With machine learning. PREVIEW- CUP. D18D20001300002).

---

**Supporting Information Fig S2.**
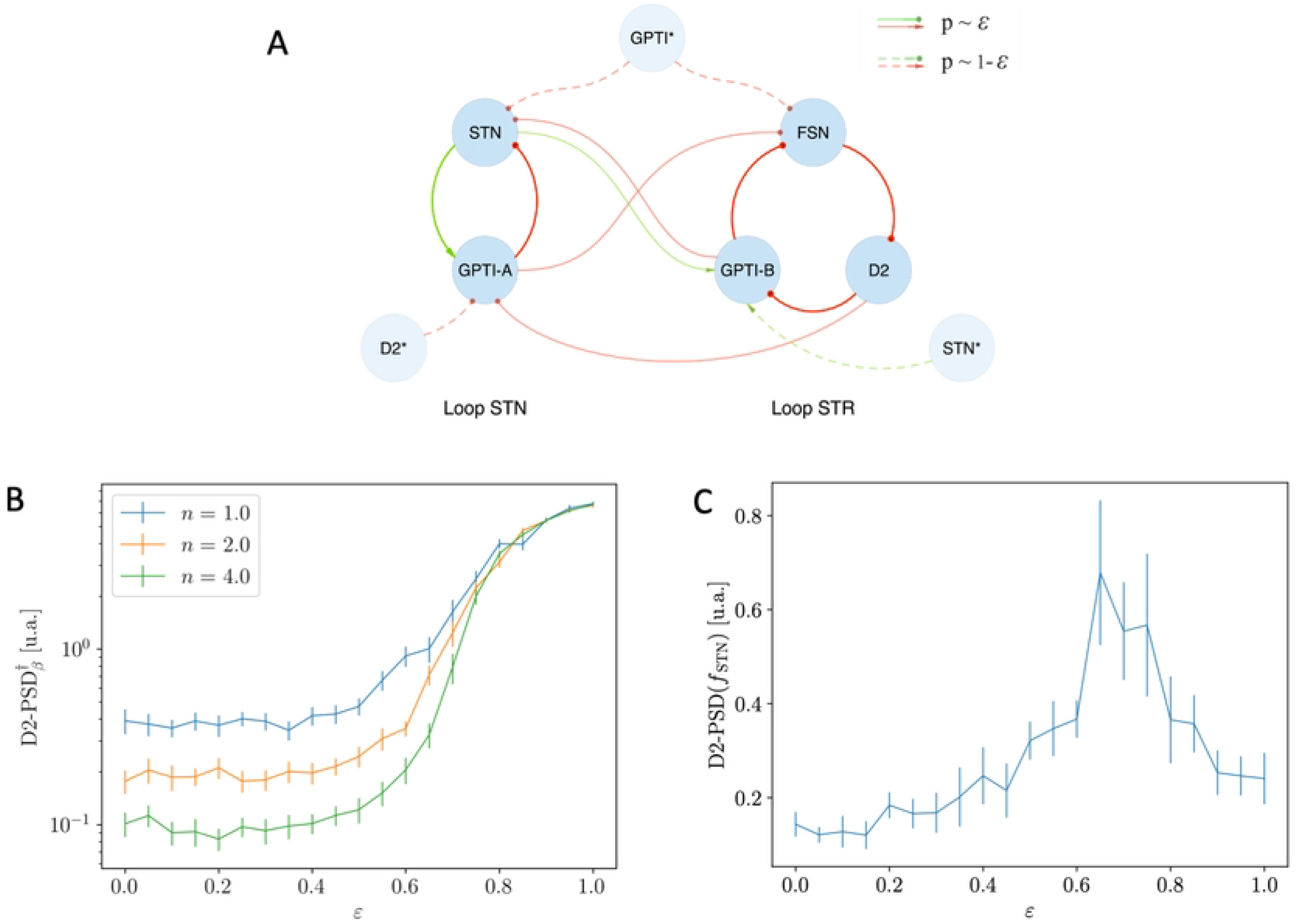

**Supporting Information Fig S3.**
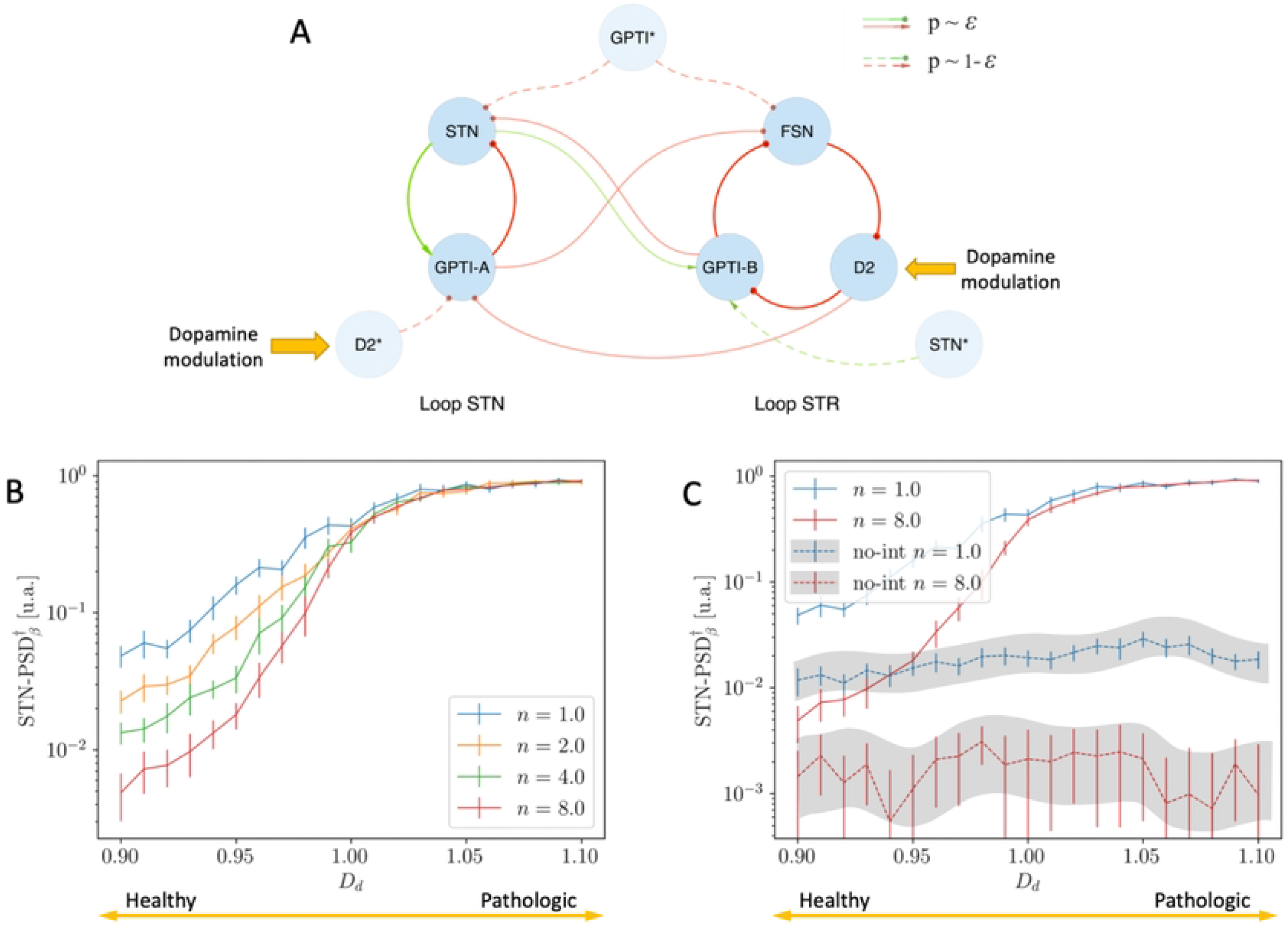

**Supporting Information Fig S4.**
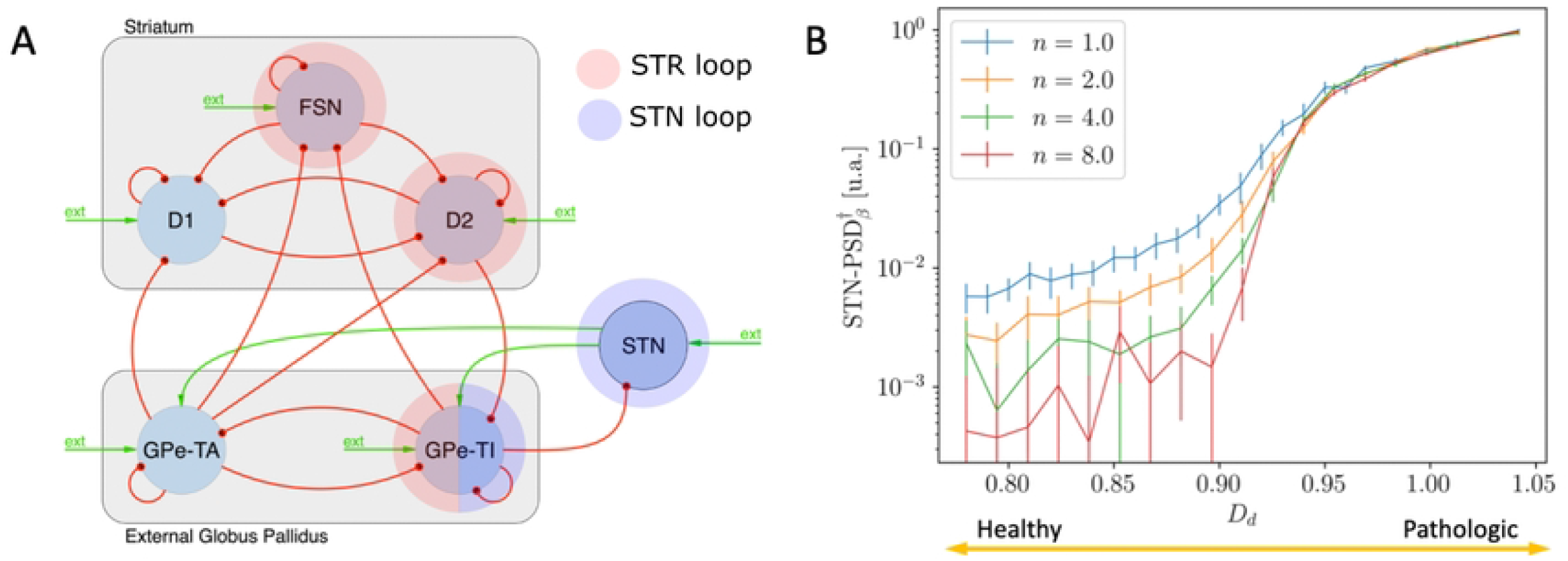

## References

1. Obeso JA, Rodriguez-Oroz MC, Stamelou M, Bhatia KP, Burn DJ. The expanding universe of disorders of the basal ganglia. The Lancet. 2014;384(9942):523–531.

2. Bergman H, Horn A. The Hidden Life of the Basal Ganglia: At the Base of Brain and Mind. Brain. 2021;.

3. Feigin VL, Nichols E, Alam T, Bannick MS, Beghi E, Blake N, et al. Global, regional, and national burden of neurological disorders, 1990–2016: a systematic analysis for the Global Burden of Disease Study 2016. The Lancet Neurology. 2019;18(5):459–480.

4. Albin RL, Young AB, Penney JB. The functional anatomy of basal ganglia disorders. Trends in neurosciences. 1989;12(10):366–375.

5. Onofrj M, Thomas A. Acute akinesia in Parkinson disease. Neurology. 2005;64(7):1162–1169.

6. Berardelli A, Rothwell JC, Thompson PD, Hallett M. Pathophysiology of bradykinesia in Parkinson’s disease. Brain. 2001;124(11):2131–2146.

7. Zimmermann R, Deuschl G, Hornig A, Schulte-Mönting J, Fuchs G, Lücking C. Tremors in Parkinson’s disease: symptom analysis and rating. Clinical neuropharmacology. 1994;.

8. Perez-Lloret S, Negre-Pages L, Damier P, Delval A, Derkinderen P, Destée A, et al. Prevalence, determinants, and effect on quality of life of freezing of gait in Parkinson disease. JAMA neurology. 2014;71(7):884–890.

9. Nutt JG, Bloem BR, Giladi N, Hallett M, Horak FB, Nieuwboer A. Freezing of gait: moving forward on a mysterious clinical phenomenon. The Lancet Neurology. 2011;10(8):734–744.

10. Nambu A. Seven problems on the basal ganglia. Current opinion in neurobiology. 2008;18(6):595–604.

11. McCarthy M, Moore-Kochlacs C, Gu X, Boyden E, Han X, Kopell N. Striatal origin of the pathologic beta oscillations in Parkinson’s disease. Proceedings of the National Academy of Sciences. 2011;108(28):11620–11625.

12. Kondabolu K, Roberts EA, Bucklin M, McCarthy MM, Kopell N, Han X. Striatal cholinergic interneurons generate beta and gamma oscillations in the corticostriatal circuit and produce motor deficits. Proceedings of the National Academy of Sciences. 2016;113(22):E3159–E3168.

13. Sharott A, Vinciati F, Nakamura KC, Magill PJ. A population of indirect pathway striatal projection neurons is selectively entrained to parkinsonian beta oscillations. Journal of Neuroscience. 2017;37(41):9977–9998.

14. Manferlotti E, Vissani M, Mazzoni A, Kumar A. Correlated inputs to striatal population drive subthalamic nucleus hyper-synchronization. In: 2021 10th International IEEE/EMBS Conference on Neural Engineering (NER). IEEE; 2021. p. 255–258.

15. Eisinger RS, Cagle JN, Opri E, Alcantara J, Cernera S, Foote KD, et al. Parkinsonian beta dynamics during rest and movement in the dorsal pallidum and subthalamic nucleus. Journal of Neuroscience. 2020;40(14):2859–2867.

16. Valsky D, Blackwell KT, Tamir I, Eitan R, Bergman H, Israel Z. Real-time machine learning classification of pallidal borders during deep brain stimulation surgery. Journal of neural engineering. 2020;17(1):016021.

17. Haumesser JK, Beck MH, Pellegrini F, Kühn J, Neumann WJ, Altschüler J, et al. Subthalamic beta oscillations correlate with dopaminergic degeneration in experimental parkinsonism. Experimental Neurology. 2021;335:113513.

18. Mallet N, Pogosyan A, Sharott A, Csicsvari J, Bolam JP, Brown P, et al. Disrupted dopamine transmission and the emergence of exaggerated beta oscillations in subthalamic nucleus and cerebral cortex. Journal of Neuroscience. 2008;28(18):4795–4806.

19. Feingold J, Gibson DJ, DePasquale B, Graybiel AM. Bursts of beta oscillation differentiate postperformance activity in the striatum and motor cortex of monkeys performing movement tasks. Proceedings of the National Academy of Sciences. 2015;112(44):13687–13692.

20. Murthy VN, Fetz EE. Oscillatory activity in sensorimotor cortex of awake monkeys: synchronization of local field potentials and relation to behavior. Journal of neurophysiology. 1996;76(6):3949–3967.

21. Plenz D, Kital ST. A basal ganglia pacemaker formed by the subthalamic nucleus and external globus pallidus. Nature. 1999;400(6745):677–682.

22. Bevan MD, Magill PJ, Terman D, Bolam JP, Wilson CJ. Move to the rhythm: oscillations in the subthalamic nucleus–external globus pallidus network. Trends in Neurosciences. 2002;25(10):525–531.

23. Sato F, Lavallée P, Lévesque M, Parent A. Single-axon tracing study of neurons of the external segment of the globus pallidus in primate. Journal of Comparative Neurology. 2000;417(1):17–31.

24. Mallet N, Pogosyan A, Márton LF, Bolam JP, Brown P, Magill PJ. Parkinsonian Beta Oscillations in the External Globus Pallidus and Their Relationship with Subthalamic Nucleus Activity. Journal of Neuroscience. 2008;28(52):14245–14258.

25. Hammond C, Bergman H, Brown P. Pathological synchronization in Parkinson’s disease: networks, models and treatments. Trends in neurosciences. 2007;30(7):357–364.

26. Alavi SM, Mirzaei A, Valizadeh A, Ebrahimpour R. Excitatory deep brain stimulation quenches beta oscillations arising in a computational model of the subthalamo-pallidal loop. Scientific reports. 2022;12(1):1–20.

27. Vitek JL, Zhang J, Hashimoto T, Russo GS, Baker KB. External pallidal stimulation improves parkinsonian motor signs and modulates neuronal activity throughout the basal ganglia thalamic network. Experimental neurology. 2012;233(1):581–586.

28. Lang AE, Zadikoff C. Parkinsonian tremor. Neurological disease and therapy. 2005;70:195.

29. Kumar A, Cardanobile S, Rotter S, Aertsen A. The role of inhibition in generating and controlling Parkinson’s disease oscillations in the basal ganglia. Frontiers in systems neuroscience. 2011;5:86.

30. Gillies A, Willshaw D, Li Z. Subthalamic–pallidal interactions are critical in determining normal and abnormal functioning of the basal ganglia. Proceedings of the Royal Society of London Series B: Biological Sciences. 2002;269(1491):545–551.

31. Pasillas-Lépine W. Delay-induced oscillations in Wilson and Cowan’s model: an analysis of the subthalamo-pallidal feedback loop in healthy and parkinsonian subjects. Biological cybernetics. 2013;107(3):289–308.

32. Merrison-Hort R, Borisyuk R. The emergence of two anti-phase oscillatory neural populations in a computational model of the Parkinsonian globus pallidus. Frontiers in computational neuroscience. 2013;7:173.

33. Holgado AJN, Terry JR, Bogacz R. Conditions for the Generation of Beta Oscillations in the Subthalamic Nucleus–Globus Pallidus Network. Journal of Neuroscience. 2010;30(37):12340–12352.

34. Holt AB, Netoff TI. Origins and suppression of oscillations in a computational model of Parkinson’s disease. Journal of computational neuroscience. 2014;37(3):505–521.

35. Leblois A, Boraud T, Meissner W, Bergman H, Hansel D. Competition between Feedback Loops Underlies Normal and Pathological Dynamics in the Basal Ganglia. Journal of Neuroscience. 2006;26(13):3567–3583.

36. van Albada SJ, Robinson PA. Mean-field modeling of the basal ganglia-thalamocortical system. I: Firing rates in healthy and parkinsonian states. Journal of Theoretical Biology. 2009;257(4):642–663.

37. van Albada SJ, Gray RT, Drysdale PM, Robinson PA. Mean-field modeling of the basal ganglia-thalamocortical system. II: dynamics of parkinsonian oscillations. Journal of theoretical biology. 2009;257(4):664–688.

38. Mallet N, Micklem BR, Henny P, Brown MT, Williams C, Bolam JP, et al. Dichotomous Organization of the External Globus Pallidus. Neuron. 2012;74(6):1075–1086.

39. Abdi A, Mallet N, Mohamed FY, Sharott A, Dodson PD, Nakamura KC, et al. Prototypic and Arkypallidal Neurons in the Dopamine-Intact External Globus Pallidus. Journal of Neuroscience. 2015;35(17):6667–6688.

40. Fujiyama F, Nakano T, Matsuda W, Furuta T, Udagawa J, Kaneko T. A single-neuron tracing study of arkypallidal and prototypic neurons in healthy rats. Brain Structure and Function. 2016;221(9):4733–4740.

41. Bevan MD, Booth PA, Eaton SA, Bolam JP. Selective innervation of neostriatal interneurons by a subclass of neuron in the globus pallidus of the rat. Journal of Neuroscience. 1998;18(22):9438–9452.

42. Crompe Bdl, Aristieta A, Leblois A, Elsherbiny S, Boraud T, Mallet NP. The globus pallidus orchestrates abnormal network dynamics in a model of Parkinsonism. Nature Communications. 2020;11(1):1570.

43. Dodson PD, Larvin JT, Duffell JM, Garas FN, Doig NM, Kessaris N, et al. Distinct Developmental Origins Manifest in the Specialized Encoding of Movement by Adult Neurons of the External Globus Pallidus. Neuron. 2015;86(2):501–513.

44. Hernández VM, Hegeman DJ, Cui Q, Kelver DA, Fiske MP, Glajch KE, et al. Parvalbumin+ Neurons and Npas1+ Neurons Are Distinct Neuron Classes in the Mouse External Globus Pallidus. Journal of Neuroscience. 2015;35(34):11830–11847.

45. Mastro KJ, Bouchard RS, Holt HA, Gittis AH. Transgenic mouse lines subdivide external segment of the globus pallidus (GPe) neurons and reveal distinct GPe output pathways. Journal of Neuroscience. 2014;34(6):2087–2099.

46. Cagnan H, Mallet N, Moll CKE, Gulberti A, Holt AB, Westphal M, et al. Temporal evolution of beta bursts in the parkinsonian cortical and basal ganglia network. Proceedings of the National Academy of Sciences. 2019;116(32):16095–16104.

47. Wilson CJ. What controls the timing of striatal spiny cell action potentials in the up state? In: The Basal Ganglia IX. Springer; 2009. p. 49–61.

48. Gage GJ, Stoetzner CR, Wiltschko AB, Berke JD. Selective activation of striatal fast-spiking interneurons during choice execution. Neuron. 2010;67(3):466–479.

49. Dong J, Hawes S, Wu J, Le W, Cai H. Connectivity and Functionality of the Globus Pallidus Externa Under Normal Conditions and Parkinson’s Disease. Frontiers in Neural Circuits. 2021;15.

50. Corbit VL, Whalen TC, Zitelli KT, Crilly SY, Rubin JE, Gittis AH. Pallidostriatal Projections Promote beta Oscillations in a Dopamine-Depleted Biophysical Network Model. Journal of Neuroscience. 2016;36(20):5556–5571.

51. Lindahl M, Kotaleski JH. Untangling basal ganglia network dynamics and function: Role of dopamine depletion and inhibition investigated in a spiking network model. Eneuro. 2016;3(6).

52. Berke JD, Okatan M, Skurski J, Eichenbaum HB. Oscillatory entrainment of striatal neurons in freely moving rats. Neuron. 2004;43(6):883–896.

53. Miller BR, Walker AG, Shah AS, Barton SJ, Rebec GV. Dysregulated information processing by medium spiny neurons in striatum of freely behaving mouse models of Huntington’s disease. Journal of neurophysiology. 2008;100(4):2205–2216.

54. Fourcaud N, Brunel N. Dynamics of the firing probability of noisy integrate-and-fire neurons. Neural computation. 2002;14(9):2057–2110.

55. Burkitt AN. A review of the integrate-and-fire neuron model: II. Inhomogeneous synaptic input and network properties. Biological cybernetics. 2006;95(2):97–112.

56. Meffin H, Burkitt AN, Grayden DB. An analytical model for the ‘large, fluctuating synaptic conductance state’typical of neocortical neurons in vivo. Journal of computational neuroscience. 2004;16(2):159–175.

57. Cavallari S, Panzeri S, Mazzoni A. Comparison of the dynamics of neural interactions between current-based and conductance-based integrate-and-fire recurrent networks. Frontiers in neural circuits. 2014;8:12.

58. Fourcaud-Trocmé N, Hansel D, Van Vreeswijk C, Brunel N. How spike generation mechanisms determine the neuronal response to fluctuating inputs. Journal of neuroscience. 2003;23(37):11628–11640.

59. Ermentrout GB, Kopell N. Parabolic bursting in an excitable system coupled with a slow oscillation. SIAM journal on applied mathematics. 1986;46(2):233–253.

60. Izhikevich EM. Simple model of spiking neurons. IEEE Transactions on neural networks. 2003;14(6):1569–1572.

61. Brette R, Gerstner W. Adaptive exponential integrate-and-fire model as an effective description of neuronal activity. Journal of neurophysiology. 2005;94(5):3637–3642.

62. Brunel N, Wang XJ. What determines the frequency of fast network oscillations with irregular neural discharges? I. Synaptic dynamics and excitation-inhibition balance. Journal of neurophysiology. 2003;90(1):415–430.

63. Oorschot DE. Total number of neurons in the neostriatal, pallidal, subthalamic, and substantia nigral nuclei of the rat basal ganglia: a stereological study using the cavalieri and optical disector methods. Journal of Comparative Neurology. 1996;366(4):580–599.

64. Hardman CD, Henderson JM, Finkelstein DI, Horne MK, Paxinos G, Halliday GM. Comparison of the basal ganglia in rats, marmosets, macaques, baboons, and humans: volume and neuronal number for the output, internal relay, and striatal modulating nuclei. Journal of Comparative Neurology. 2002;445(3):238–255.

65. Butcher J. Practical Runge–Kutta methods for scientific computation. The ANZIAM Journal. 2009;50(3):333–342.

66. O’Neill ME. PCG: A family of simple fast space-efficient statistically good algorithms for random number generation. ACM Transactions on Mathematical Software. 2014;.

67. West TO, Magill PJ, Sharott A, Litvak V, Farmer SF, Cagnan H. Stimulating at the right time to recover network states in a model of the cortico-basal ganglia-thalamic circuit. PLoS computational biology. 2022;18(3):e1009887.

68. Suryanarayana SM, Hellgren Kotaleski J, Grillner S, Gurney KN. Roles for globus pallidus externa revealed in a computational model of action selection in the basal ganglia. Neural Networks. 2019;109:113–136.

69. Blenkinsop A, Anderson S, Gurney K. Frequency and function in the basal ganglia: the origins of beta and gamma band activity. The Journal of Physiology. 2017;595(13):4525–4548.

70. Chakravarty K, Roy S, Sinha A, Nambu A, Chiken S, Hellgren Kotaleski J, et al. Transient Response of Basal Ganglia Network in Healthy and Low-Dopamine State. eNeuro. 2022;9(2):ENEURO.0376–21.2022.

71. Tachibana Y, Iwamuro H, Kita H, Takada M, Nambu A. Subthalamo-pallidal interactions underlying parkinsonian neuronal oscillations in the primate basal ganglia. European Journal of Neuroscience. 2011;34(9):1470–1484.

72. Pavlides A, Hogan SJ, Bogacz R. Computational models describing possible mechanisms for generation of excessive beta oscillations in Parkinson’s disease. PLoS computational biology. 2015;11(12):e1004609.

73. Rubin JE. Computational models of basal ganglia dysfunction: the dynamics is in the details. Current opinion in neurobiology. 2017;46:127–135.

74. Lai HY, Liao LD, Lin CT, Hsu JH, He X, Chen YY, et al. Design, simulation and experimental validation of a novel flexible neural probe for deep brain stimulation and multichannel recording. Journal of neural engineering. 2012;9(3):036001.

75. Butenko K, Bahls C, Schröder M, Köhling R, van Rienen U. OSS-DBS: Open-source simulation platform for deep brain stimulation with a comprehensive automated modeling. PLoS computational biology. 2020;16(7):e1008023.

76. Fleming JE, Dunn E, Lowery MM. Simulation of closed-loop deep brain stimulation control schemes for suppression of pathological beta oscillations in Parkinson’s disease. Frontiers in neuroscience. 2020;14:166.

77. Tinkhauser G. The present and future role of clinical neurophysiology for Deep Brain Stimulation; 2022.

78. Shah A, Nguyen TAK, Peterman K, Khawaldeh S, Debove I, Shah SA, et al. Combining multimodal biomarkers to guide deep brain stimulation programming in Parkinson disease. Neuromodulation: technology at the neural interface. 2022;.

79. Tinkhauser G, Moraud EM. Controlling Clinical States Governed by Different Temporal Dynamics With Closed-Loop Deep Brain Stimulation: A Principled Framework. Frontiers in neuroscience. 2021;15.

